# Proteomics reveals how the tardigrade damage suppressor protein teaches transfected human cells to survive UV-C stress

**DOI:** 10.1101/2023.06.19.545547

**Authors:** Enxhi Shaba, Claudia Landi, Carlotta Marzocchi, Lorenza Vantaggiato, Luca Bini, Claudia Ricci, Silvia Cantara

## Abstract

The genome sequencing of the tardigrade *Ramazzottius varieornatus* revealed a unique nucleosome-binding protein, named Damage Suppressor (Dsup), which resulted to be crucial for the extraordinary abilities of tardigrades in surviving extreme stresses, such as UV. Evidence in Dsup transfected human cells, suggests that Dsup mediates an overall response of DNA damage signaling, DNA repair and cell cycle regulation resulting in an acquired resistance to stress. Given these promising outcomes, our study attempts to provide a wider comprehension of the molecular mechanisms modulated by Dsup in human cells, and to explore the Dsup-activated molecular pathways under stress. We performed a differential proteomic analysis of Dsup-transfected and control human cells, under basal condition and at 24-hour recovery after exposure to UV-C. We demonstrate by enrichment and network analyses, for the first time, that even in the absence of external stimuli and more significantly after stress, Dsup activates mechanisms involved with the Unfolded Protein Response, the mRNA processing and stability, cytoplasmic stress granules, the DNA Damage Response and the telomere maintenance. In conclusion, our results shed new light on Dsup-mediated protective mechanisms, and increase our knowledge of the molecular machineries of extraordinary protection against UV-C stress.

## 1. Introduction

The Damage Suppressor (Dsup) is a unique nucleosome-binding protein discovered for the first time in 2016 when scientists completed the sequencing of the *Ramazzottius varieornatus* tardigrade genome [1]. Around 1,464 species of tardigrades are found worldwide to date [2], and they are considered aquatic because they require a layer of water around their bodies to prevent dehydration. Tardigrades have been observed from the deep sea to sand dunes and they are able to survive extreme environments, including space. In particular, X-ray protection seems to be exerted by Dsup thanks to a direct bound to chromatin [3], but tardigrades can also tolerate increased temperatures and pressure [4,5] as well as dehydration [6]. Due to these amazing properties, Dsup has gained great interest among the scientific community. In particular, human and plant cells have been engineered to produce Dsup, resulting in the acquisition of a higher resistance to X-rays [1], oxidative stress [3] and genomutagens [7]. Recently, it has been demonstrated that Dsup affects the expression of endogenous genes under stress conditions such as UV-C exposure. Particularly, mRNA evidence showed that the expression of genes involved in DNA repair and cell cycle checkpoints is up-regulated together with transcription factors modulation [8]. According to their extraordinary potentiality, tardigrades may represent a great chance of exploring new ways of adaption to surrounding environments. In fact, as commonly accepted, our planet is facing massive changes, principally due to human abuse and unlimited exploitation of the Earth’s resources. These events are leading to the extensive alteration of macro- and micro-environments, such as drought and weather changes, therefore the exposure to these phenomena is often unbearable for common living organisms, from plants to humans [9]. In particular, given the progressive disruption of the atmospheric ozone layer, the understanding of UV radiation’s impact on living organisms and the investigation of potential protective solutions became extremely urgent. UV radiation might be classified into three primary types: ultraviolet A (UV-A, 315-399 nm), ultraviolet B (UV-B, 280-314 nm), and ultraviolet C (UV-C, 100-279 nm). UV-A are not filtered by the ozone layer, whereas UV-B are only absorbed partially. Even if UV-C are completely absorbed by the ozone layer and atmosphere, they represent the most damaging radiation type as the lower the wavelength, the higher the energy content of UV-radiation. Especially, they cause cyclobutane pyrimidine dimers (CPDs) and pyrimidine-6,4-pyrimidinone photoproducts (6,4PPs) on exposed DNA, as well as UV-associated ROS production by photodynamic reactions exerting high DNA damaging effects [10]. Both CPDs and 6,4PPs lesions alter DNA structure, introducing bends that inhibit cell transcription and replication. After the replicative arrest, cells undergo a progressive telomere erosion, which contributes to genetic instability and accelerates cell death by apoptosis [11,12]. In addition, telomeres are hypersensitive to single-strand DNA damage, therefore their protection is essential for cell life and DNA stability. Given these increasing damaging risks, the strong necessity of exploring new ways of adaptation to these newly forming environmental scenarios, such as tardigrades’ do, suits perfectly with high-throughput technologies such as OMIC sciences. In fact, these approaches are able to provide a wider perspective and a more comprehensive knowledge of biological complexity at multi-levels.

To this purpose, we applied a functional proteomic approach investigating the molecular mechanisms modulated by Dsup in transfected human cells, and exploring the Dsup-activated molecular pathways under UV-C stress. In particular, we performed a differential proteomic analysis of Dsup-transfected (Dsup+) and control (Dsup−) HEK293T cells, under basal condition and at 24-hour recovery after 15 seconds exposure to UV-C. Various molecular pathways and differential proteins of interest were tested to confirm their modulation in presence of Dsup at cellular level.

## 2. Results

### 2.1. In the absence of external stimuli, Dsup-transfected cells present a modulation of proteins involved in protein folding, telomere maintenance and metabolic processes

In order to understand whether Dsup itself was able to activate cellular mechanisms in the absence of external stresses, we applied a differential proteomic analysis between Dsup− and Dsup+ cells at basal conditions. Like that, we highlighted 29 differentially abundant proteins (Table S1 and in Figure S1), of which 19 were identified by MALDI-ToF. As reported by heatmap analysis in Figure 1A, 17 were higher abundant and 12 were lower abundant after Dsup transfection. In particular, supervised PCA (Figure 1B) shows that Dsup− and Dsup+ proteomic profiles well separate from each other, highlighting the Dsup impact in the cellular proteome. Based on PCA, the increased abundance of eukaryotic initiation factor 4A-II (IF4A2), actin, cytoplasmic 1 (ACTB), heat shock 70 kDa protein 1A N-term fragment (HS71A_HS71C N-frag) and the decreased abundance of asparagine synthetase [glutamine-hydrolyzing] (ASNS), heterogeneous nuclear ribonucleoproteins A2/B1 (ROA2), peroxiredoxin 1 (PRDX1), T-complex protein 1 subunit delta (TCPD) and tyrosine-tRNA ligase, cytoplasmic (SYYC), have a greater relevance in Dsup−/+ cell culture discrimination.

**Figure 1.**
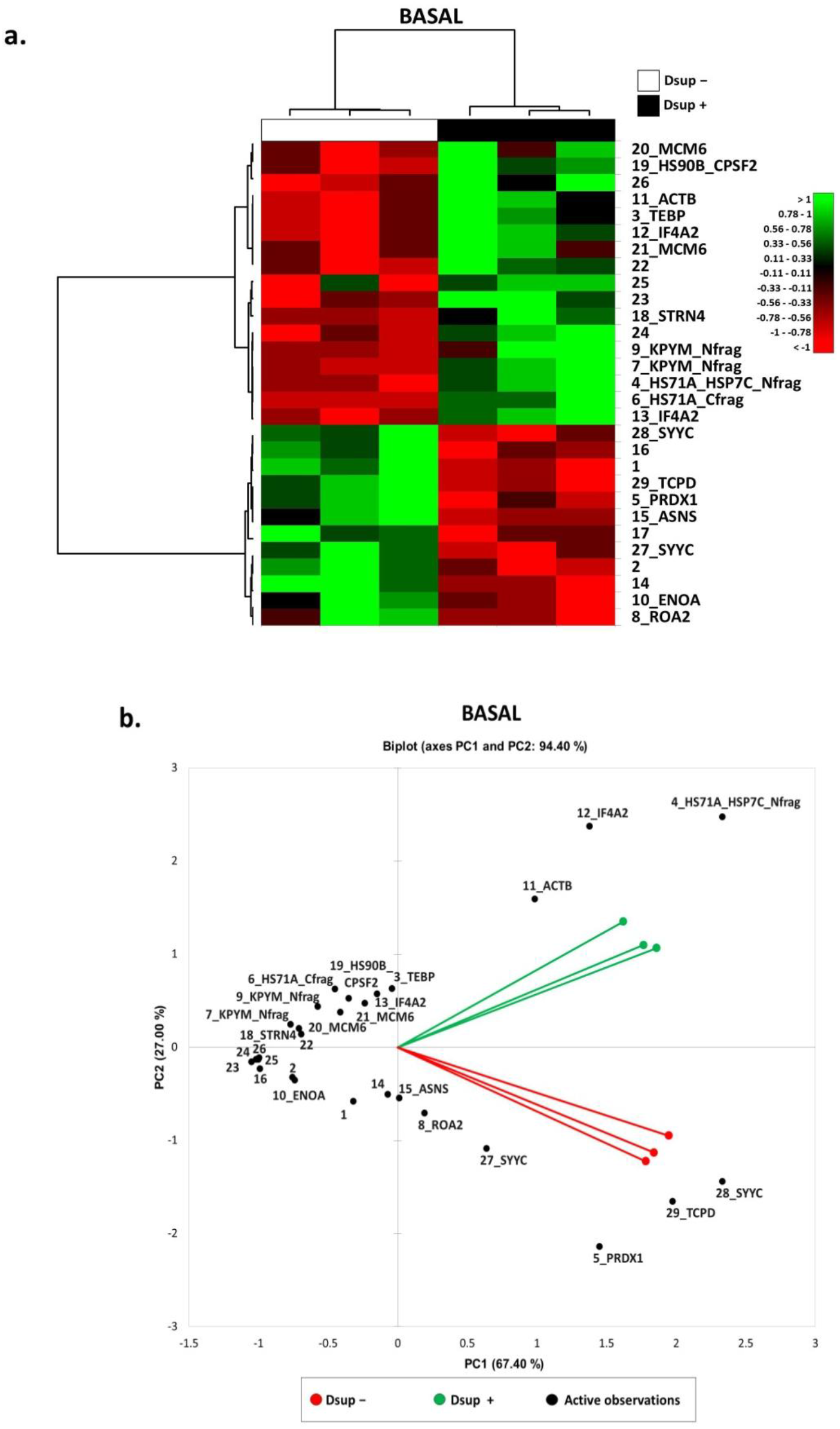
a) Heatmap analysis of the differential spots found by proteomic analysis of Dsup− vs Dsup+ HEK293T cell cultures at basal condition. As shown in the legendhigh abundant spots are reported in green, while low abundant ones are reported in red. Black numbers and acronyms represent differential protein spots, while Dsup− cell cultures are reported in black and Dsup+ cell cultures are reported in white. b) Principal Component Analysis of the differential spots found by proteomic analysis of Dsup− vs Dsup+ HEK293T cell cultures at basal condition. PCA summarizes a 94% of variance (PC1: 67.40%; PC2: 27%). Black numbers and acronyms represent differential protein spots, while Dsup− cell cultures are reported in red and Dsup+ cell cultures are reported in green.

Figure 2A shows the protein network of the differentially higher abundant proteins in Dsup+ basal cells with HSP70, HSP90 and PKM2 acting as central functional hubs, suggesting their pivotal role in down-stream or up-stream molecular pathways regulation following Dsup transfection. On the other hand, Figure 2B reports the protein network of the differentially lower abundant proteins in basal cells after Dsup transfection, showing that ENO, ASNS, ROA2, PRDX1 and TCPD are in this case central functional hubs. In addition, an enrichment analysis of GO Biological Processes of all differential proteins in Dsup−/+ cells was performed (Figure 2C). Our results highlight that the differential proteome of Dsup+ cells is characterized by a modulation of proteins involved in chaperone-mediated protein complex assembly (HSP90, p23 co-chaperone, HSP70), protein folding and refolding (TCP1-delta, peroxiredoxin, HSP90, p23 co-chaperone, HSP70, HSC70), telomere maintenance and organization (hnRNP A2, HSP90, p23 co-chaperone, HSP70), ATP metabolic process (ENO, HSP70, PKM2, HSC70, Pyruvate kinase), cellular nitrogen compound metabolic process, (ENO, hnRNP A2, eIF4A, HSP90, p23 co-chaperone, HSP70, PKM2, HSC70, MCM6, CPSF2, ASNS) and regulation of cellular response to stress (ENO, Peroxiredoxin, HSP90, p23 co-chaperone, HSP70, HSC70).

**Figure 2.**
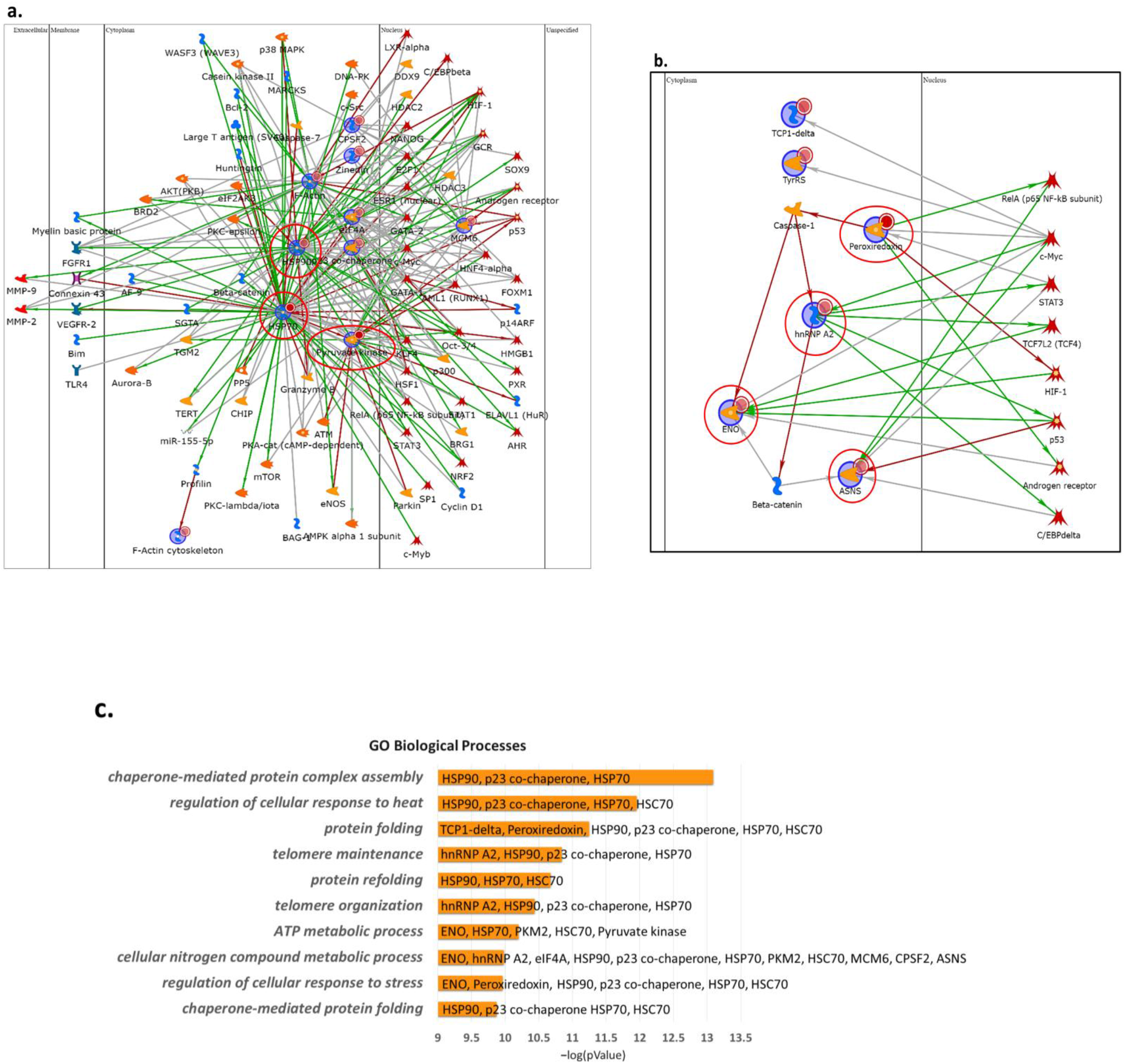
Network analysis by MetaCore software to highlight characteristic protein interactions of the higher- (a) and lower-abundant (b) differential protein species of Dsup− vs Dsup+ HEK293T cell cultures analysis at basal condition. a) HSC70, HSP90 beta, PKM2 and HSP90 are the central functional hubs of the net and are marked in red circles. b) ENO1, ASNS, ROA2, PRDX1 and TCPD are the central functional hubs, marked in red circles. c) Enrichment analysis by GO Biological Processes by MetaCore software: each histogram represents an enriched GO Biological Process associated to its p-value in −log10 and its related differential proteins

Moreover, enrichment analysis by map folders (Figure 3) evidenced transcription regulation and DNA damage response as two important molecular mechanisms regulated by identified proteins. In particular, transcription regulation is related to the negative regulation of HIF1α function and HIF-1 targets (HSP90, HSP70, HSC70, ENO, PKM2), in addition to mTORC1 downstream signalling (eIF4A), whereas DNA damage response is related to the regulation of telomere length (HSP90, TEBP/p23 co-chaperone), apoptosis and survival by granzyme A signaling (ROA2, HSP70), ATM and ATR (Figure 3).

**Figure 3.**
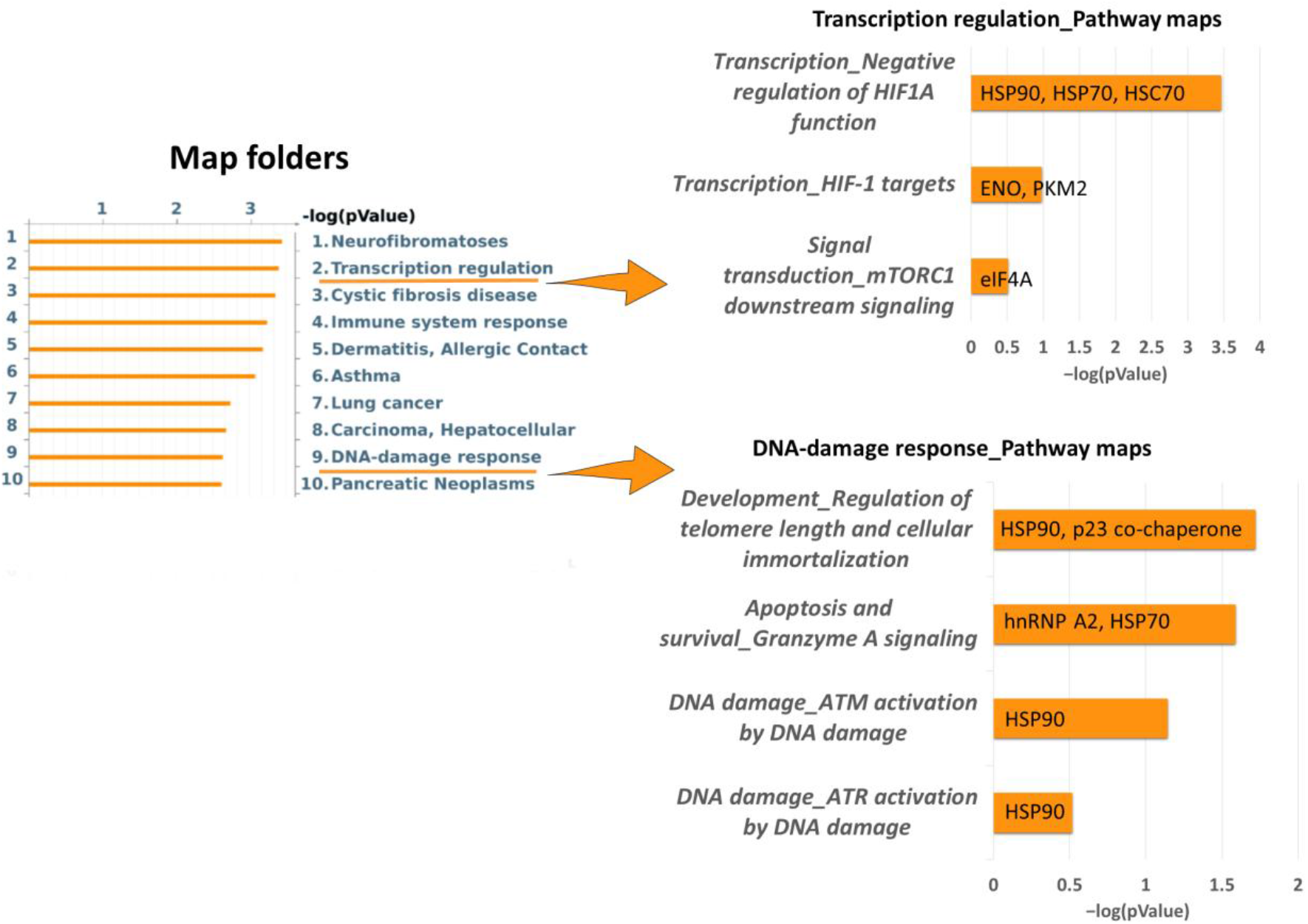
Enrichment analysis by map folders analysis by MetaCore software, performed with all differential proteins of Dsup− vs Dsup+ proteomic analysis (basal condition). Figure also reports Pathway Maps in Folder ‘Transcription regulation’ and pathway Maps in Folder ‘DNA-damage response’. Each histogram represents an enriched map folder/pathway map associated to its p-value in −log10 and its related differential proteins

### 2.2 Proteins involved in DNA damage response and regulation of transcription are increased in Dsup+ cells after 15” of UV-C irradiation followed by 24h of recovery

Our group already demonstrated that Dsup+ cells exposed UV-C and allowed to recover for 24h presented a statistically significant increase in survival rate compared to Dsup− (Figure S2) with the activation of transcription factors and overexpression of ATM and ATR mRNAs [8]. Considering these previous data, we aimed to investigate deeper which cellular mechanisms might be activated by Dsup to promote cell survival after UV-C radiation at protein level. Similarly, to what performed in basal conditions, we performed a differential proteomic analysis of Dsup−/+ cells exposed to UV-C. We highlighted 63 differentially abundant protein spots (Table S2 and Figure S3), of which 46 were identified by MALDI-ToF. The heatmap in Figure 4A shows that 46 proteins were differentially higher abundant, while 17 proteins were differentially lower abundant after Dsup transfection. Moreover, Figure 4B reports the PCA of Dsup−/+ cells in a 2D spatial distribution, showing that Dsup significantly impacts on the differential proteome after UV-C irradiation, with a higher extent compared to basal conditions. Particularly, it also highlighted that the decreased amount of PPIA, PCBP1, ESTD, IDH3A, PIPNB, ARPC5, THIO, LTOR2/RFA3 and EI2BB is correlated to the distribution of Dsup− cells, whereas the increased amount of HS90B, ROA2, SERA, PRPS1, HS71A/B, HNRPU, PRDX1 and TIF1B to Dsup+ cells.

**Figure 4.**
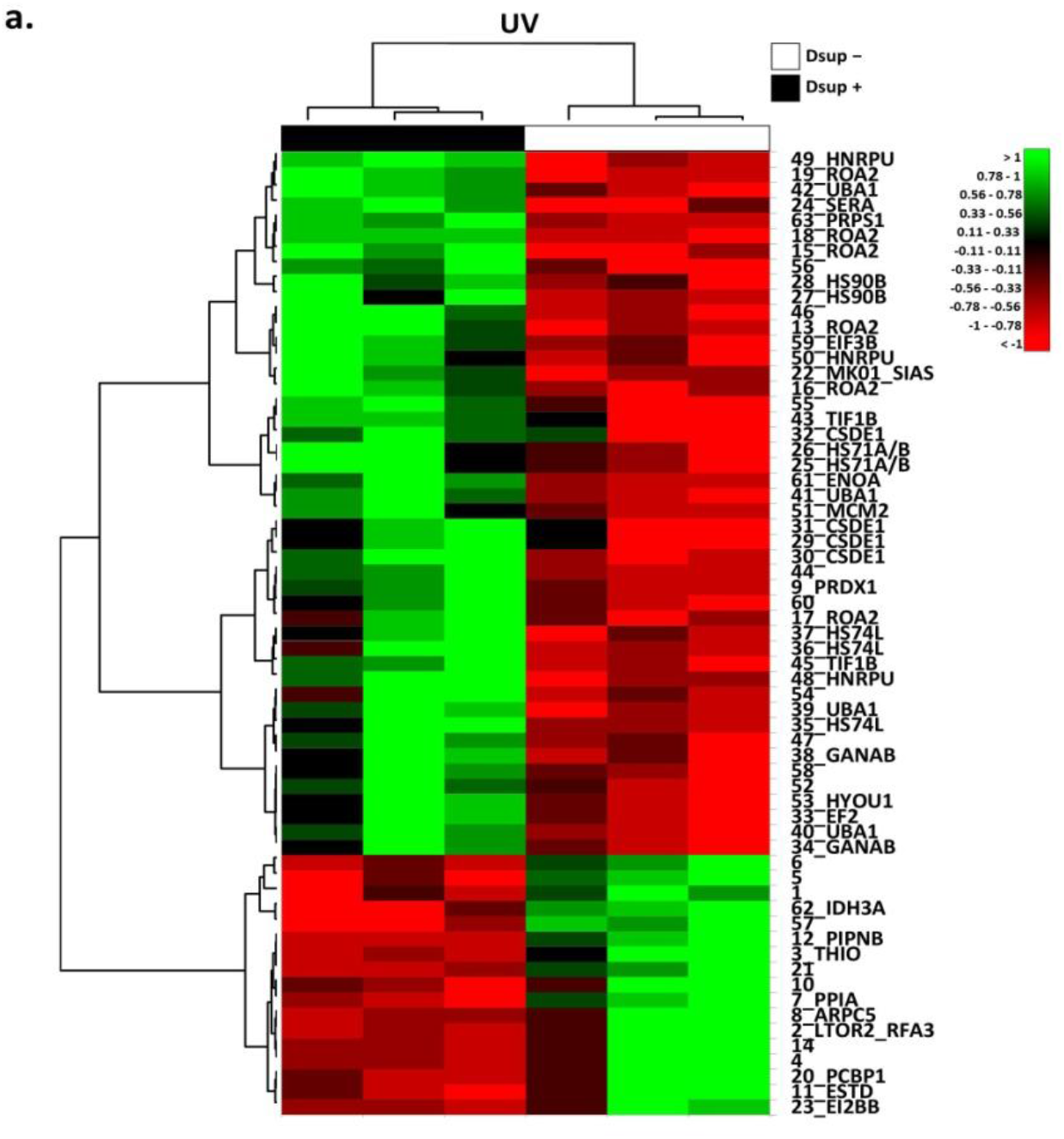
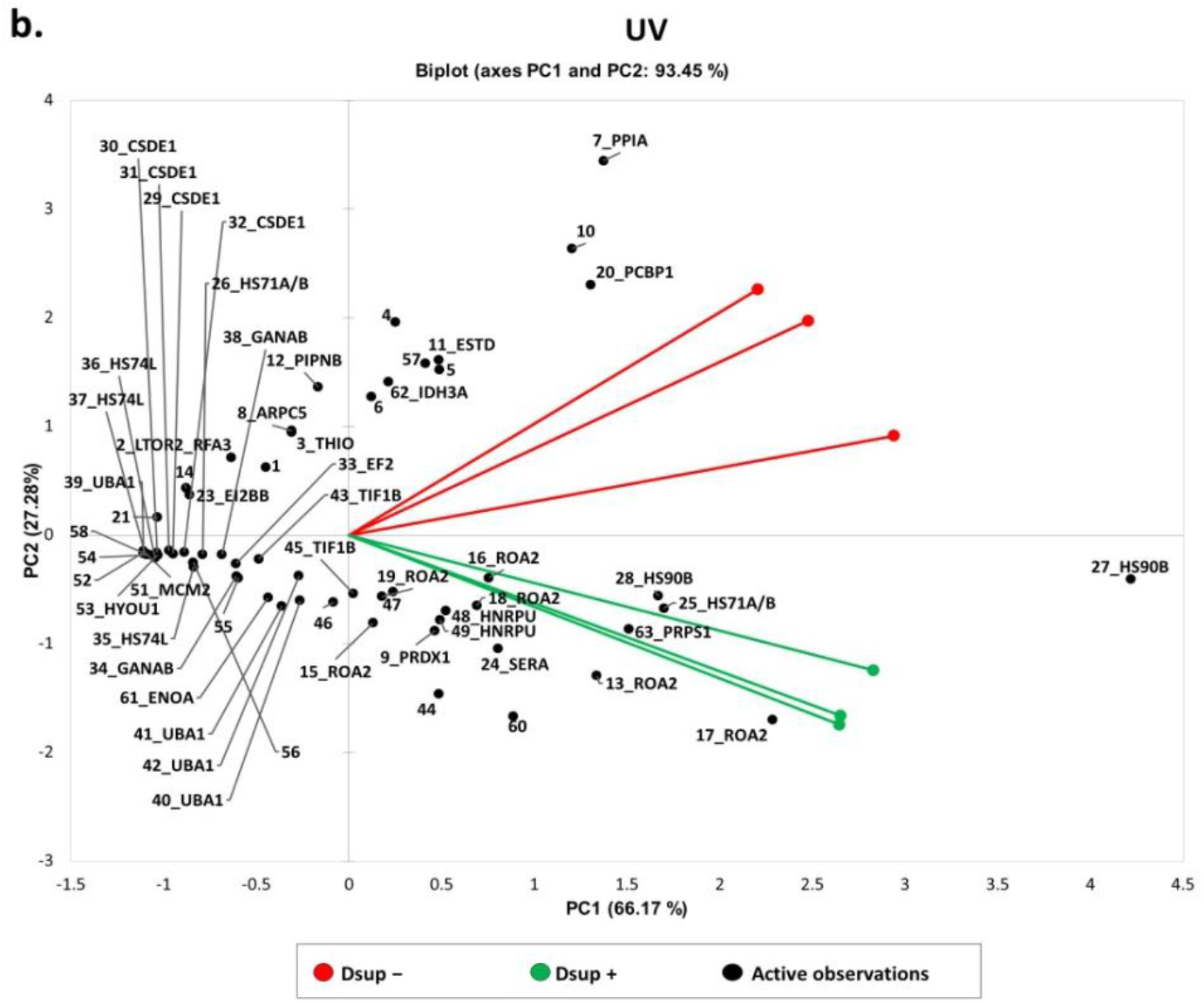
**(a)** Heatmap analysis of the differential spots found by proteomic analysis of Dsup− vs Dsup+ HEK293T cell cultures at 24h recovery after 15 seconds of UV-C irradiation. As shown in the legend, high abundant spots are reported in green, while low abundant ones are reported in red. Black numbers and acronyms represent differential protein spots, while Dsup− cell cultures are reported in white and Dsup+ cell cultures are reported in black. **(b)** Principal Component Analysis of the differential spots found by proteomic analysis of Dsup− vs Dsup+ HEK293T cell cultures at 24h recovery after 15 seconds of UV-C irradiation. PCA summarize 93.45% of variance subdivided for 66.17% in PC1 and 27.28% in PC2. Black numbers and acronyms represent differential protein spots, while Dsup− cell cultures are reported in red and Dsup+ cell cultures are reported in green.

An enrichment analysis by map folders of differentially higher abundant proteins in Dsup+ cells is reported in Figure 5A, showing three relevant molecular pathways, each grouping particular mechanisms associated with the identified higher abundant proteins: transcription regulation, protein degradation and DNA damage response. Figure 5A.I shows specific pathways related to transcription regulation: we put in evidence the higher abundance of two protein species of HSP90 (spots 27 and 28) and of HSP70 (spot 25 and 26), together with the higher amount of DNA replication licensing factor (MCM2; spot 51), implicated with a negative regulation of HIF1A. We also found the higher abundance of 4 proteoforms of ubiquitin-like modifier-activating enzyme 1 (UBA1; spot 39-42), as well as the involvement of two protein species of transcription intermediary factor 1-beta (TIF1B; TIF1-beta; spots 43, 45), involved in NF-kB signaling-related processes and in the role of heterochromatin protein 1 family in transcriptional silencing, respectively. Secondly, Figure 5A.II reports additional pathways associated with protein degradation, such as ubiquitin-proteasomal pathways, suggested by the higher amount of HSP70 and UBA1, as well as signaling transduction processes, advised by the more abundant mitogen-activated protein kinase 1 (MK01; ERK1/2; spot 22). Furthermore, Figure 5A.III evidences particular pathways related to the DNA damage response (DDR), such as ATM/ATR activation by DNA damage and regulation of telomere length and cellular immortalization associated with the higher amount of HSP90 and MK01.

**Figure 5.**
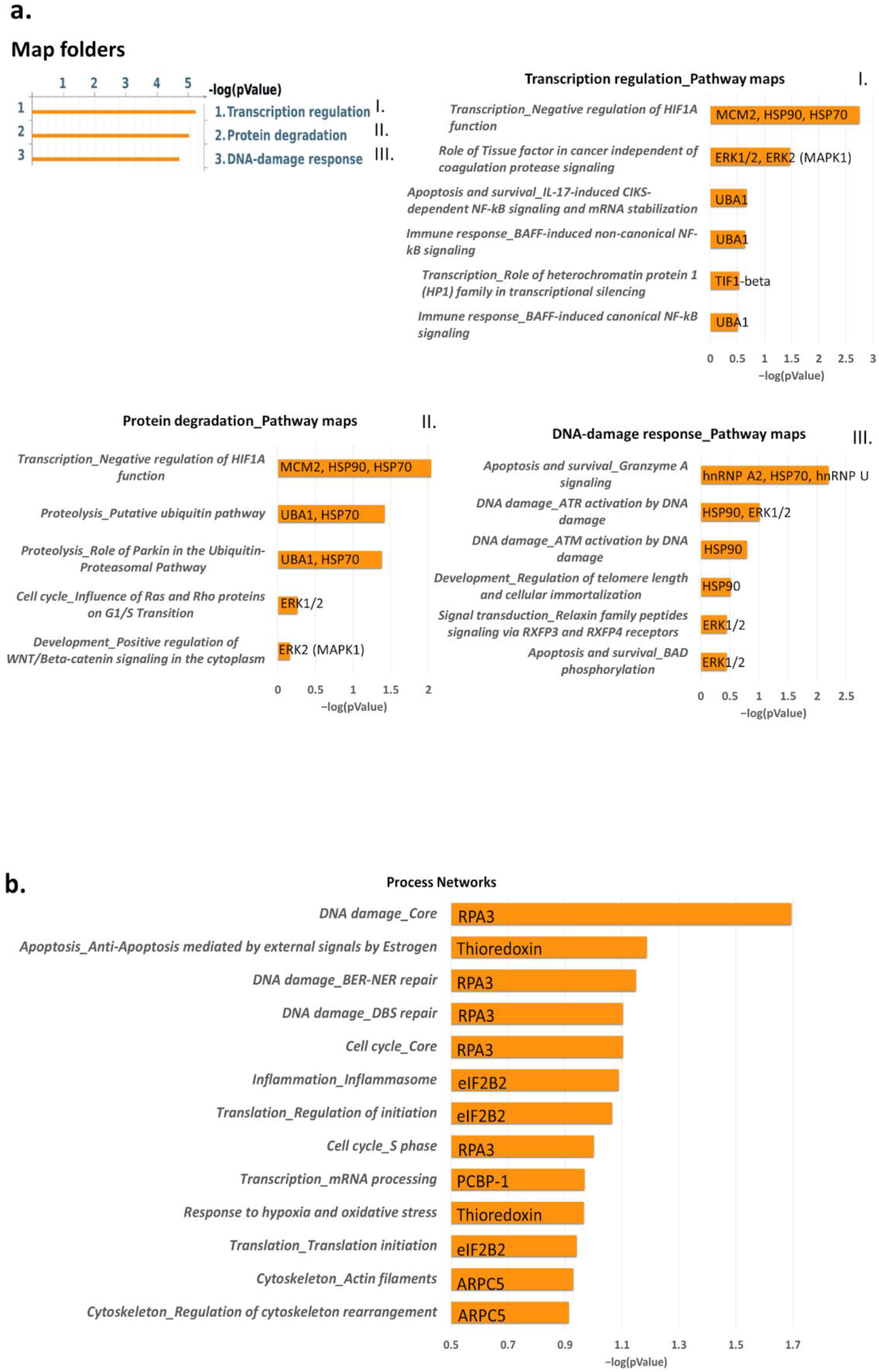
**a)** Enrichment analysis by map folders analysis by MetaCore software, performed with the differentially higher abundant proteins in Dsup+ HEK293T cells at 24h recovery after 15 seconds of UV-C irradiation. Figure also reports Pathway Maps in Folder ‘Transcription regulation’ (I), in Folder ‘Protein degradation’ (II) and in Folder ‘DNA-damage response’ (III). **b)** Enrichment analysis by process networks analysis by MetaCore software, performed with the differentially low abundant proteins in Dsup+ HEK293T cells at 24h recovery after 15 seconds of UV-C irradiation. Each histogram represents an enriched map folder/pathway map/process network associated to its p-value in −log10 and its related differential proteins.

On the other side, an enrichment analysis by process networks of differentially lower abundant proteins in Dsup+ cells is reported in Figure 5B. Particularly, DNA damage response processes, including base excision repair (BER)/nucleotide excision repair (NER) events and double-strand DNA breaks (DSB) repair, as well as cell cycle regulatory processes are reported in association with the replication protein A 14 kDa subunit (RFA3; RPA3; spot 2). Transcriptional regulatory mechanisms of mRNA processing and translation are associated with the lower abundance of the translation initiation factor eIF2B subunit beta (EI2BB; eIF2B2; spot 23). Other relevant pathways are the response to hypoxia and oxidative stress (THIO; spot 3), as well as the regulation of cytoskeleton rearrangement indicated by the lower amount of actin-related protein 2/3 complex subunit 5 (ARPC5; spot 8).

In order to explore potential interactions among the identified proteins, a network analysis was performed (Figure 6), where TIF1-beta, ERK1/2, ENO1, Thioredoxin and MCM2 were reported to be central functional hubs. In particular, TIF1-beta showed several interactions with other differentially abundant proteins, such as peptidyl-prolyl cis-trans isomerase A (PPIA; cyclophilin A; spot 7), eukaryotic initiation factor 4AII (IF4A2; eIF4A; spot 12, 13), eukaryotic translation initiation factor 3 subunit B (EIF3B; eIF3; spot 59) and elongation factor 2 (EF2; eEF2; spot 33). In addition, MCM2 displayed interactions with ATR serine/threonine protein kinase, HIF-1 and c-Myc, which was reported to be connected to heterogeneous nuclear ribonucleoproteins A2/B1 (ROA2; hnRNP A2; spot 13, 15-19) and U (hnRNP U; spot 48-50), as well as cold shock domain-containing protein E1 (CSDE1; UNR; spot 29-32).

**Figure 6.**
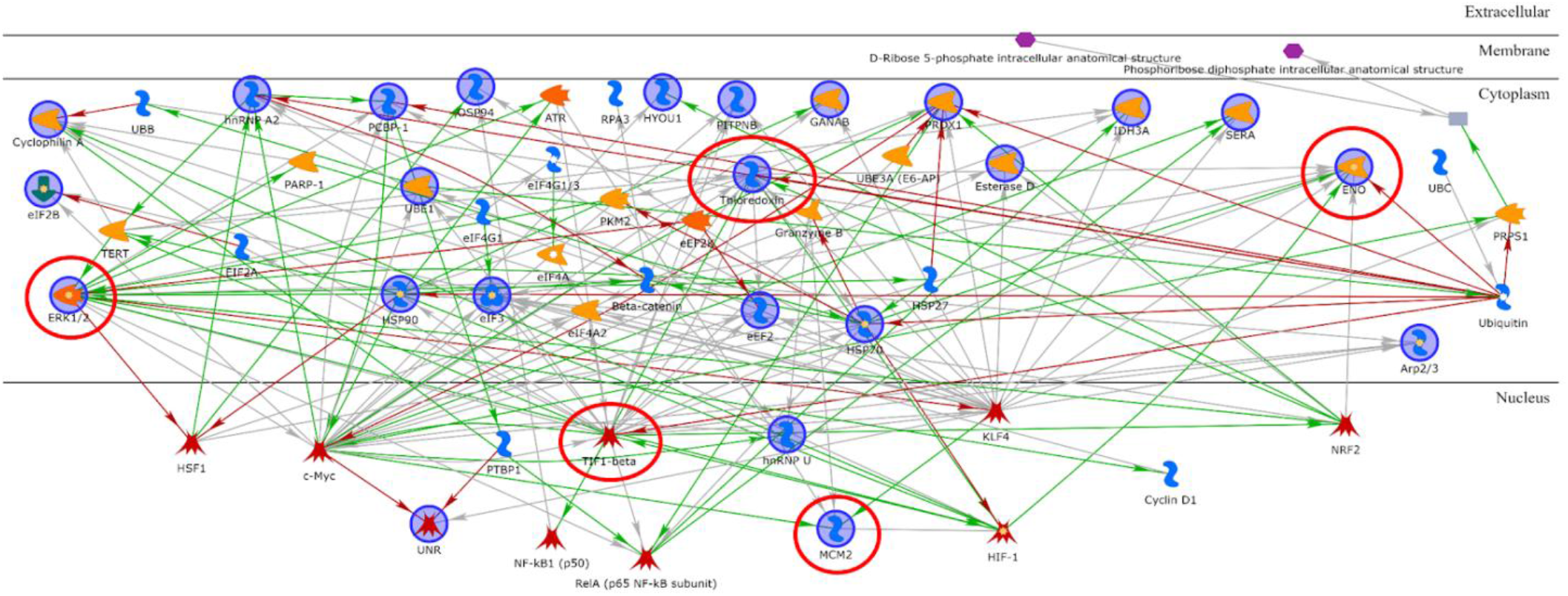
Network analysis by MetaCore software to highlight characteristics protein interactions of the all differential protein species of the proteomic analysis of Dsup− vs Dsup+ HEK293T cell cultures at 24h recovery after 15 seconds of UV-C irradiation. Thioredoxin, ERK1/2, TIF1-beta, MCM2 and ENOA are the central functional hubs of the net and are marked in red circles.

### 2.3 Targeted validation of proteins belonging to Unfolded Protein Response (UPR), transcription and metabolic regulation, DDR and telomere length regulation pathways

Given the differential proteomic data and the specific molecular pathways suggested by bioinformatic analysis, a targeted validation was performed. HSP90 is a differential protein found higher abundant in Dsup+ cells in basal conditions and after UV-C irradiation. As a central factor in protein folding processes, as well as transcription regulation, DNA repair and regulation of metabolic processes following cell stress [13], Dsup-mediated HSP90 increased abundance might be a crucial effector of Dsup protection. As shown in Figure 7A, western blot analysis confirms a moderate increase in abundance in Dsup+ cells at basal level. Interestingly, HSP90 abundance in Dsup− cells drastically decreases after UV-C irradiation, while HSP90 levels remain almost unchanged between basal and irradiated conditions in Dsup+ cells, thus showing a significant increase after UV-C irradiation, compared to the Dsup− counterpart.

**Figure 7.**
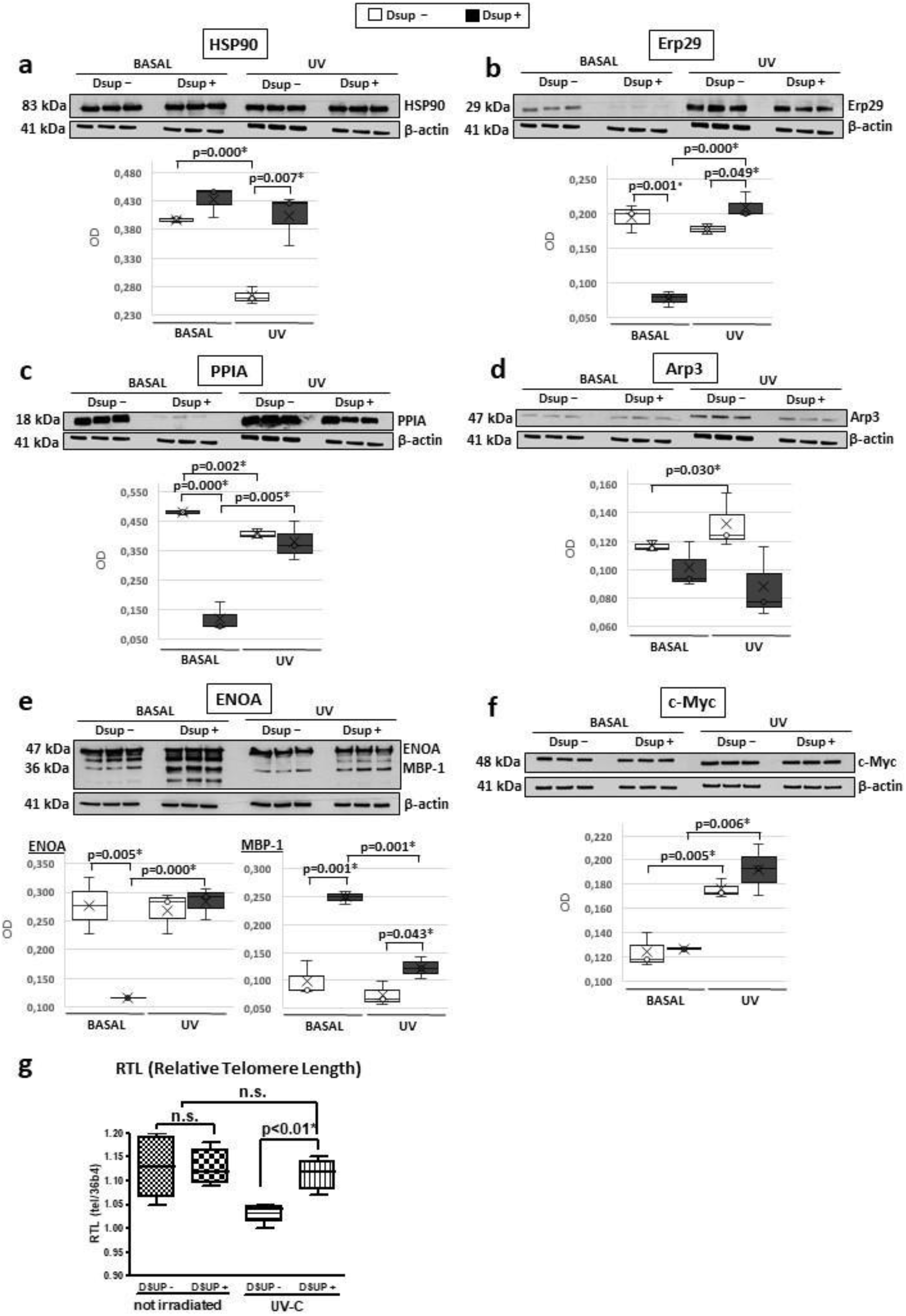
Western blot validations and Relative Telomere Length (RTL) assay. Figure reports WB validations of HSP90 **(a)**, Erp29 **(b)**, PPIA **(c)**, Arp3 **(d)**, ENOA and MBP-1 **(e)**, c-Myc **(f)** and normalization of bands’ intensity was performed on total β-actin. Legend displays that normalized WB lane intensities of Dsup–cells are reported as white, while those of Dsup+ cells are reported in black.Differences are considered significant with p-value <0.05. Relative Telomere Length (RTL) assay **(g)** displays the measurement by real time PCR of the relative telomere length in Dsup− cells compared to Dsup+ cells exposed to UV-C. P<0.01 by one-way ANOVA with Fisher’s correction.

Since UPR is a molecular process highlighted by our proteomic results, we evaluated Erp29 protein levels. Erp29 abundance, indeed, increases in response to endoplasmic reticulum stress (ERS) [14], especially in the case of genotoxic stress induced by ionizing radiation [15]. As showed in Figure 7B, Erp29 significantly decreases in Dsup+ cells at basal level, and then statistically increases after UV-C irradiation in comparison to Dsup− cells. Indeed, Dsup− cells show a decreased level of Erp29 after UV-C treatment.

Since PPIA is a multifunctional protein that acts also in UPR, we performed a western Blot assay which highlights a significant decreased abundance of PPIA in Dsup+ basal cells, in addition to a significant decrease in Dsup− cells and a significant increase in Dsup+ cells after UV-C irradiation compared to their basal counterpart (Figure 7C). ARPC5, that we found lower abundant in Dsup+ cells after UV-C stress, is a subunit of the Arp2/3 complex [16]. Therefore, we decided to perform a western blot assay for Arp3, to verify if another member of the complex is modulated by Dsup transfection, since the complex promotes actin polymerization in the nucleus regulating gene transcription and the DNA damage repair [17]. As expected, also Arp3 is lower abundant in Dsup+ cells at basal level and after UV-C stress, with respect to the corresponding Dsup− cells, displaying the same trend of ARPC5 and suggesting that the entire complex is low abundant in Dsup+ cells (Figure 7D).

Notably, proteomic data also suggest a cellular metabolic modulation in Dsup+ cells, as indicated by the dysregulation of different protein species of ENOA. Figure 7E highlights an interesting trend of ENOA proteoforms validated by western blot. In basal conditions, Dsup+ cells present various proteoforms at lower molecular weight than the referring protein at 47 kDa, which results significantly increased in abundance in Dsup+ cells, confirming proteomic data. There is a particular ENOA proteoform (37kDa), referred to as c-Myc Binding Protein 1 (MBP-1) [18] and missing of the first 90 amminoacids, which appears to be significantly higher abundant in Dsup+ basal and UV-C irradiated cells, compared to the Dsup− counterparts. This proteoform is reported to inhibit the expression of c-Myc, which is in turn statistically increased at protein level in both Dsup− and Dsup+ cells after UV-C stress (with a higher extent for Dsup+ cells, Figure 7F).

Since proteomics highlights proteins involved in telomere maintenance, we investigated telomere stability in Dsup− and Dsup+ cells after UV-C irradiation by measuring the relative telomere length (RTL) by qPCR. As shown in Figure 7G, RTL is significantly longer in Dsup+ cells compared to Dsup− cells at 24h recovery after UV-C exposure (p<0.01). No differences in RTL were found in non-irradiated cells and between non irradiated cells and Dsup+ cells after UV-C exposure (Figure 7G).

## 3. Discussion

Dsup protein was reported for the first time by Hashimoto T. et al as a prominent example of tardigrade-unique abundant proteins involved in tolerability, as it resulted to be a DNA-associating protein mediating DNA protection against radiation in cultured animal and human cells. (Hashimoto, 2015). For these reasons, our study focused on applying a functional proteomic approach, aimed to investigate Dsup impact in human cells in basal conditions and after UV-C exposure, and to investigate the potentially modulated mechanisms in response to induced stress.

In this study we evidenced for the first time that, even at basal conditions, Dsup protein impacts on cells by modulating proteins associated with the UPR. Indeed, at basal conditions, we highlighted several differentially abundant proteins belonging to protein folding, telomere maintenance and metabolic processes pathways. One of the main players we found to be highly abundant in Dsup+ human cells is HSP90, which is reported to significantly mediate cellular resistance via UPR pathway [20]. Our results also report the differential abundance of ENOA, which is reported to act as a glycolytic enzyme, a plasminogen receptor and a DNA binding protein [24]. Moreover, it is reported to be involved in protective processes by enhancing cell viability and glycolysis against various stresses [25–28]. In our study, ENOA activation was particularly evident after UV-C exposure. Interestingly, ENOA presents several proteoforms that are related to multiple functions and significance [28]. Of notice, we detected an increased amount of a 37kDa proteoform in Dsup+ cells, which could be the myc promoter-binding protein 1 (MBP-1), a shorter protein variant of ENOA involved in the c-myc regulation [18] and reported to be increased as a consequence of cell response to stress conditions [29].

As a result of UV exposure, there is the formation of helix-disturbing lesions, such as bulky DNA adducts (CPDs and 6,4PPs), which interfere with DNA transcription and replication, leading also to the production of double-strand breaks (DSBs) [30]. DSBs might also occur in UV-C irradiated cells as a consequence of single-strand breaks (SSBs) repair failure by Base Excision Repair (BER) [31,32]. In addition, UV light might also indirectly cause DNA damage via ROS production in photodynamic reactions leading to ROS-damaged bases and direct SSBs [10]. The occurrence of these lesions canonically triggers the DDR via ATR and/or ATM activation, which initiates signaling pathways of DNA damage sensing and repair. The main activated DNA repair mechanism following UV-induced DNA damages is Nucleotide Excision Repair (NER) [33]. Interestingly, our results indicate a higher abundance of one protein species of ERK1/2 in Dsup+ UV-C stressed cells, and especially ERK1/2 is associated to DDR signaling pathway by enrichment analysis. Indeed, the activation of ERK1/2 has been reported to be activated by UV exposure [34] and to potentially modulate Nucleotide Excision Repair (NER) pathways [35]. Moreover, our functional proteomic analysis reports a higher abundance of two protein species of TIF1B (TIF1B; TRIM28; KAP1) in Dsup+ UV-C stressed cells, which is reported to be involved in DNA damage repair mechanisms, also following UV exposure [36,37]. The network analysis suggests that TIF1B represents a functional hub interacting with several differential proteins in Dsup+ UV-C stressed cells. Among these, it regulates the transcription of ARPC5 (subunit of the Arp2/3 complex) [38], which supports actin polymerization in the nucleus, contributing to gene transcription and DNA repair processes [39]. In Dsup+ cells we observed a concomitant lower abundance of Arp3 and the replication factor A 14 kDa subunit (RPA3), the latter being the smallest subunit of RPA complex which is an essential coordinator of DNA repair [40]. Moreover, network analysis also reports that TIF1B transcriptionally regulates UBA1 [41] and we observed the higher abundance of 4 proteoforms of UBA1 in Dsup+ cells after UV-C exposure. Indeed, the ubiquitination pathway has a positive regulatory role for efficient NER machinery, potentially recruiting repair factors to DNA damage sites and UBA1 is reported to be potentially involved in the repair of UV-induced DNA damage [42]. In addition, network analysis also suggests that in Dsup+ UV-C stressed cells TIF1B interacts with telomerase reverse transcriptase (TERT), as previously reported by Agarwal N. et al., who describes that TIF1B phosphorylation by mTORC1 allows it to induce TERT transcription [43]. The telomere maintenance mechanisms might be crucial events in Dsup+ cells, as supported by the dysregulation of several other differential proteins, such as Prostaglandin E synthase 3 (TEBP/p23 co-chaperone), HSP90, HNRPU and ROA2 [44–47]. Our functional analysis further suggests a potential interaction between TERT and PPIA [48], which is decreased in abundance in Dsup+ cells exposed to UV-C and implicated in the regulation of cellular processes, such as protein folding, transcriptional regulation, cell survival and response to stress [49]. Of interest, Daneri-Becerra et al. report PPIA to be a mitochondrial factor which complexes with HSP90 and p23 co-chaperone, playing a significant role in cell survival upon stress [50]. Moreover, PPIA is also reported as functional interacting partner of TAR DNA binding protein (TARDBP or TDP-43), which is predominantly an RNA/DNA binding protein involved in RNA processing and metabolism, including RNA transcription, splicing, transport and stability. Specifically, TARDBP is a key component of hnRNP complexes, in which hnRNP A2/B1 is the major heterogeneous ribonucleoprotein recognized by PPIA, essential in regulating the maturation of newly formed nuclear RNAs/pre-mRNAs into messenger RNAs (mRNAs) [51–54]. Interestingly, in Dsup+ cells exposed to UV-C, we report the increased abundance of numerous proteoforms of heterogeneous nuclear ribonucleoproteins A2/B1 (ROA2), in addition to heterogeneous nuclear ribonucleoprotein U (hnRPU). In light of these findings, Dsup seems to strongly impact on the modulation of mRNA stability processes, as also suggested by the down regulation in Dsup+ cells of Poly(rC)-binding protein 1 (PCBP1), a specific RNA-binding protein that associates with cytoplasmic polyadenylation elements contributing to mRNA stability and translational activity [55].

Cellular responses to environmental stresses often target a fine regulation of the transcriptome by activating a set of conserved processes aimed to restore cellular homeostasis, side by side with the UPR. One crucial event is represented by the formation of membraneless compartments, referred to as cytoplasmic stress granules (SGs), especially composed of untranslated mRNAs, translation initiation factors, small ribosomal subunits and RNA-binding proteins (RBPs). Their main advantages are the reduced energy consumption, the regulation of proteostasis and the cell survival improvement under damaging conditions [56]. Many differentially higher abundant proteins in Dsup+ cells are associated with SGs, such as the 4 proteoforms of CSDE1 [57,58], EIF3B, IF4A2 and ROA2. In details, IF4A2 (component of the eIF4F complex) is required for the so-called “non-canonical” stress granule formation that, among its components, include also EIF3B (component of the eIF3 complex) and ROA2 (as part of the required hnRNPs) [59]. Of notice, this stress response is only transient, leading SGs to be disassembled when the insult is removed and the translation is restored. Recently, SGs definition has being evolving into dynamic cytoplasmic biocondensates, in which intrinsically disordered proteins (IDPs) are essential for the assembly [60]. IDPs are characterized by a lack of a persistent tertiary and/or secondary structure, making them extremely dynamic and flexible [61]. Dsup has been recognized as a IDP [62]. The potential involvement of IDPs in the formation of other protective structures, like SGs, in response to diverse stresses might broaden the Dsup-mediated protective impact in human cells.

## Conclusions

Our study shows that in transfected human cells, Dsup mediates cell protection against the damaging effects of UV-C by activating more efficient mechanisms of DNA damage repair, mRNA stability, telomere elongation and maintenance, unfolded protein response and cytoplasmic stress granules response together with a metabolic modulation. These promising data are, however, just a little part of the still unknown cellular pathways involved in Dsup-mediated protection from external stimulus, therefore investigation must proceed forward.

## 4. Materials and Methods

### 4.1 Cell cultures, Dsup transfection and UV-C radiation

The HEK293T cell line was kindly donated by Prof. Sandra Donnini (University of Siena). The HEK293T cells (mycoplasma-free, verified by N-GARDE Mycoplasma PCR reagent set, Euroclone) were maintained in Dulbecco’s modified Eagle’s Medium (DMEM) supplemented with 10% fetal bovine serum (FBS). pCXN2KS-Dsup was a gift from Prof. Kunieda Takekazu (Addgene plasmid #90019; http://n2t.net/addgene.90019; RRID: Addgene_90019). Empty vector (same plasmid without Dsup) was kindly donated from Dr. Jlenia Brunetti (University of Siena). The expression construct and the empty plasmid were transfected into HEK293T cells using Lipofectamine® 2000 Reagent (Life Technologies), and stably transfected cells were selected by 700 µg/ml G418 (SERVA Electrophoresis GmbH) treatment for 3 weeks. The evaluation of the Dsup transcript presence was assessed as described in previous work [8] and shown in Figure S4.

In order to evaluate Dsup-induced resistance against radiation, both Dsup+ and Dsup− cells were plated at 100,000 cells/mL density in a 96-well plate. Before treatment, complete medium was removed and replaced with100 µL PBS and cells were exposed to UV-C for 15” (source 8W lamp - 4 mJ/cm2, wavelength 260-280 nm, source distance 5 cm). After treatment, cells were incubated in complete medium for 24h and then recovered [8].

### 4.2 Proteomic analysis

Three independent cell cultures for basal and treatment conditions, in both cells transfected with Dsup (Dsup+) and empty vector (Dsup−), were harvested by trypsinization and washed three times with phosphate buffer saline (PBS), with brief centrifugation (5 min at 1600 × g) after each wash. Cell pellets were stored at −80 °C until proteomic analysis. Samples were dissolved in lysis buffer (7 M Urea, 2 M Thiourea, 4% w/v 3-[(3-cholamidopropyl) dimethylammonia]-1-propanesulfonate hydrate (CHAPS) and 1% w/v dithioerythritol (DTE)) and protein concentration was determined by Bradford assay. Samples were resolved by two-dimensional electrophoresis (2DE), according to Puglia M et al [64]. Image analysis was carried out Melanie™ Classic 9 (SIB Swiss Institute of Bioinformatics, Geneva, Swiss) and the percentage of relative volume (%V) of each spot was exported used for statistical analysis. Statistically significant differences by ANOVA test were determined by XLStat (Addinsoft, Paris, France) and processed according to the ratio value ≥ 2 of corresponding %Vol means.

### 4.3 Protein identification by MALDI-ToF mass spectrometry

Differential proteins found were identified by MALDI-ToF mass spectrometry by Peptide Mass Fingerprint (PMF). Differential spots were manually excised from MS-compatible silver stained gels. Spots were destained first in a solution of 30 mM potassium ferricyanide and 100 mM sodium sulphate anhydrous, later in 200 mM ammonium bicarbonate. Then, dehydrated in 100% acetonitrile (ACN). Protein spots were rehydrated and digested overnight at 37 °C in trypsin solution. Digested proteins were then placed on MALDI target, dried and covered with matrix solution of 5 mg/ml α-cyano-4-hydroxycinnamic acid (CHCA) in 50% v/v ACN and 0.5% v/v trifluoroacetic acid (TFA). The MS analysis was carried out using the UltrafleXtreme™ MALDI-ToF/ToF mass spectrometer (Bruker Daltoniks, Bremen, Germany) equipped with a 200 Hz smartbeam™ I laser in the positive reflector mode with the following parameters: 80 ns of delay; ion source 1: 25 kV; ion source 2: 21.75 kV; lens voltage: 9.50 kV; reflector voltage: 26.30 kV and reflector 2 voltage: 14.00 kV. The applied laser wavelength and frequency were 353 nm and 100 Hz, respectively, and the percentage was set to 50%. Final mass spectra were produced by averaging 1500 laser shots targeting five different positions within the spot. MS spectra were acquired and processed by the FleXanalysis software version 3.0 (Bruker), using peptides arising from trypsin autoproteolysis as internal standard for calibration. The resulting mass lists were filtered for common contaminants, such as matrix-related ions, trypsin autolysis and keratin peaks. Protein identification was carried out by Peptide Mass Fingerprinting search using MASCOT (Matrix Science Ltd., London, UK, http://www.matrixscience.com; accessed on 02.05.2022), setting up the following parameters: Homo sapiens as taxonomy, SwissProt as database, 20 ppm as mass tolerance, one admissible missed cleavage site, carbamidomethylation (iodoacetamide alkylation) of cysteine as fixed modification and oxidation of methionine as a variable modification. The mass spectrometry proteomics data have been deposited to the ProteomeXchange Consortium via the PRIDE [65] partner repository with the dataset identifier PXD037439.

### 4.4 Telomere length

Relative telomere length (RTL) was quantified by qPCR on live cells. Briefly, culture medium was removed from cell cultures and the adhering, living cells were washed with PBS to remove possible debris and death cells. Once PBS was removed, adhering cells were harvested by trypsinization and DNA was extracted by QiAmp DNA mini kit (Qiagen) following kit instructions. RTL was quantified on 100 ng/µl of DNA by determining the relative ratio of telomere (T) repeat copy number to a single copy gene (S) copy number (T/S ratio). This ratio is proportional to the average telomere length [66]. 36B4, encoding acidic ribosomal phosphoprotein P0, was used as the single copy gene. Primers were as follow and were used at 300 nM final concentration in a reaction mix containing 12.5 µl SYBER GREEN PCR Master Mix for a final volume of 25 µl.: telomere sense, 5’-GGTTTTTTGAGGGTGAGGGTGAGGGTGAGGGTGAGGGT-3’; telomere antisense, 5’-TCCCCGACTATCCCTATCCCTATCCCTATCCCTATCCCTA-3’; 36B4 sense, 5’-CCCATTCTATCACAACGGTACAA-3’, 36B4 antisense, 5’-CAGCAAGTGGGAAGGTGTAATCC-3’. The thermal cycling profile for the telomere amplification was: 95° for 10 min followed by 30 cycles of 95°C for 15 s and 54°C for 1 min; for the 36B4 amplification was: 95° for 10 min followed by 40 cycles of 95°C for 15 s and 60°C for 1 min. To exclude the presence of nonspecific binding between SYBER GREEN and primers, a melting curve was added at the end of all PCR amplification reactions.

### 4.5 Western Blot

Twentyfive µg of each protein sample were suspended in 20 µl of LAEMMLI (Tris HCl pH 6.8 62.5 mM, 20% v/v glycerol, 2% w/v SDS, 5% v/v β-mercaptoethanol, 0.032% w/v bromophenol blue), heated at 95°C for 7 min and loaded in 12% polyacrylamide gels. Monodimensional electrophoresis was carried out using the Mini-PROTEAN electrophoresis system (Bio-Rad). After SDS-PAGE, gels were equilibrated for 1 h in transfer buffer and then transferred to nitrocellulose membranes (Cytiva). After transfer, membranes were stained by Red Ponceau (0.2% w/v Ponceau S in 3% acetic acid) to assess the correct and equal protein loading, as well as correct protein transfer. Membranes were tested with different primary antibodies, such as anti-HSP90 (mouse monoclonal, 610418, BD Transduction Laboratories, New Jersey, U.S., dilution 1:1000), anti-ERP29 (rabbit polyclonal, ALX-210-404-R100, Enzo Life Sciences, NY, U.S., dilution 1:2500), anti-ENOA (mouse monoclonal, sc-100812 Santa Cruz biotechnology, Dallas, U.S., dilution 1:200), anti c-Myc (rabbit polyclonal, ab69987, Abcam, Cambridge, UK, dilution 1:1000), anti-PPIA (rabbit polyclonal, LS-C36063, LifeSpan Biosciences, Seattle, U.S., dilution 1:1000), anti-ARP3 (rabbit polyclonal, sc-15390, Santa Cruz Biotechnology, Dallas, U.S., dilution 1:200). Membranes were incubated with different secondary antibodies, according to specific antibody composition, such as peroxidase-conjugated goat anti-rabbit at 1:7000 dilution and peroxidase-conjugated goat anti-mouse at 1:3000 dilutions.

Considering that, by proteomic analysis, only one proteoform of β-actin in Dsup−/+ cells at basal condition varies in abundance, western blot normalization against total β-actin in full was performed (anti-β-actin mouse monoclonal, A5441, Sigma, U.S., dilution: 1:40000). ECL chemiluminescence detection system (Cytiva) was utilized to specific immunoreactive proteins and images were analysed with ImageJ (1.51p, NIH, Chicago, IL, USA) to obtain band intensity. Statistical analysis was performed by XLSTAT (Addinsoft).

### 4.6 Statistical and enrichment analyses

Multivariate analysis by Principal Component analysis (PCA) and heatmap analysis were performed for basal and UV-C treatment, comparing Dsup− and Dsup+ cells, considering the %V of statistically significant differential spots by XLStat (Addinsoft). Enrichment analysis was performed by submitting the UniProt accession numbers (AN) of identified proteins to the MetaCore software (https://portal.genego.com, date access 23.05.2022) (Clarivate analytics, Boston, MA, USA). For Dsup+ vs. Dsup− comparison, the protein network analysis was then carried out by the “shortest path” algorithm that builds a hypothetical network connecting two experimental proteins directly or indirectly using one or more MetaCore database proteins. This algorithm allows building a network including only closely related proteins and introducing a maximum of one non experimental protein prioritized according to their statistical significance (p≤0.001). Networks were visualized graphically as nodes (proteins) and edges (links between proteins) and relevant biological processes were then prioritized according to their statistical significance (p≤0.001 and FDR) reporting the specific proteins involved (legend is fully explained in Figure S5). GO Biological Processes enrichment analysis was also performed. Moreover, Map Folder tool was used to highlight well-known molecular pathway regulation processes including our differential proteins, as well as enrichment by process networks and pathway maps. For telomere length, statistical analysis was performed with StatView for Windows, version 5.00.1 (SAS Institute, Cary, NC). One-way ANOVA with Fisher’s correction as used to determine RTL difference as a continuous variable. Survival data were analyzed using a paired test for non-parametric data (Wilcoxon signed-rank test). For all comparisons, a p value of <0.05 was considered significant.

## Supplementary Materials

Figure S1: Representative 2DE electropherograms of Dsup− vs Dsup+ HEK293T cells at basal condition; Figure S2: Survival assay of HEK293T cells control (Dsup-) and Dsup-transfected (Dsup+) after 15″ of UV-C exposure recovered after 24h. The dotted line represents the basal condition (100%); Figure S3: Representative 2DE electropherograms of Dsup− vs Dsup+ HEK293T cells exposed to 15 seconds of UV-C radiation and recovered after 24h; Figure S4: Dsup assessment. Dsup expression in control (Dsup−) and transfected (Dsup+) HEK293T cells was performed by endpoint PCR and visualized in 2% agarose gel by ethidium bromide; Figure S5: MetaCore legend; Table S1: Proteomic results of the Dsup− vs Dsup+ HEK293T cells analysis at basal condition (DOC); Table S2: Proteomic result of the Dsup− vs Dsup+ HEK293T cells exposed to UV-C analysis (DOC).

