## supplementary figures for "Proteomics reveals how the tardigrade damage suppressor protein teaches transfected human cells to survive UV-C stress"

**Supporting Information:** This article contains supporting information, as listed below:

**Figure S1:** Representative 2DE electropherograms of Dsup− vs Dsup+ HEK293T cells at basal condition;

**Figure S2:** Survival assay of HEK293T cells control (Dsup-) and Dsup-transfected (Dsup+) after 15” of UV-C exposure recovered after 24h. The dotted line represents the basal condition (100%)

**Figure S3:** Representative 2DE electropherograms of Dsup− vs Dsup+ HEK293T cells exposed to 15 seconds of UV-C radiation and recovered after 24h;

**Figure S4:** Dsup assessment. Dsup expression in control (Dsup−) and transfected (Dsup+) HEK293T cells was performed by endpoint PCR and visualized in 2% agarose gel by ethidium bromide;

**Figure S5:** MetaCore legend;

**Table S1**: Proteomic results of the Dsup− vs Dsup+ HEK293T cells analysis at basal condition

**Table S2**: Proteomic result of the Dsup− vs Dsup+ HEK293T cells analysis exposed to UV-C

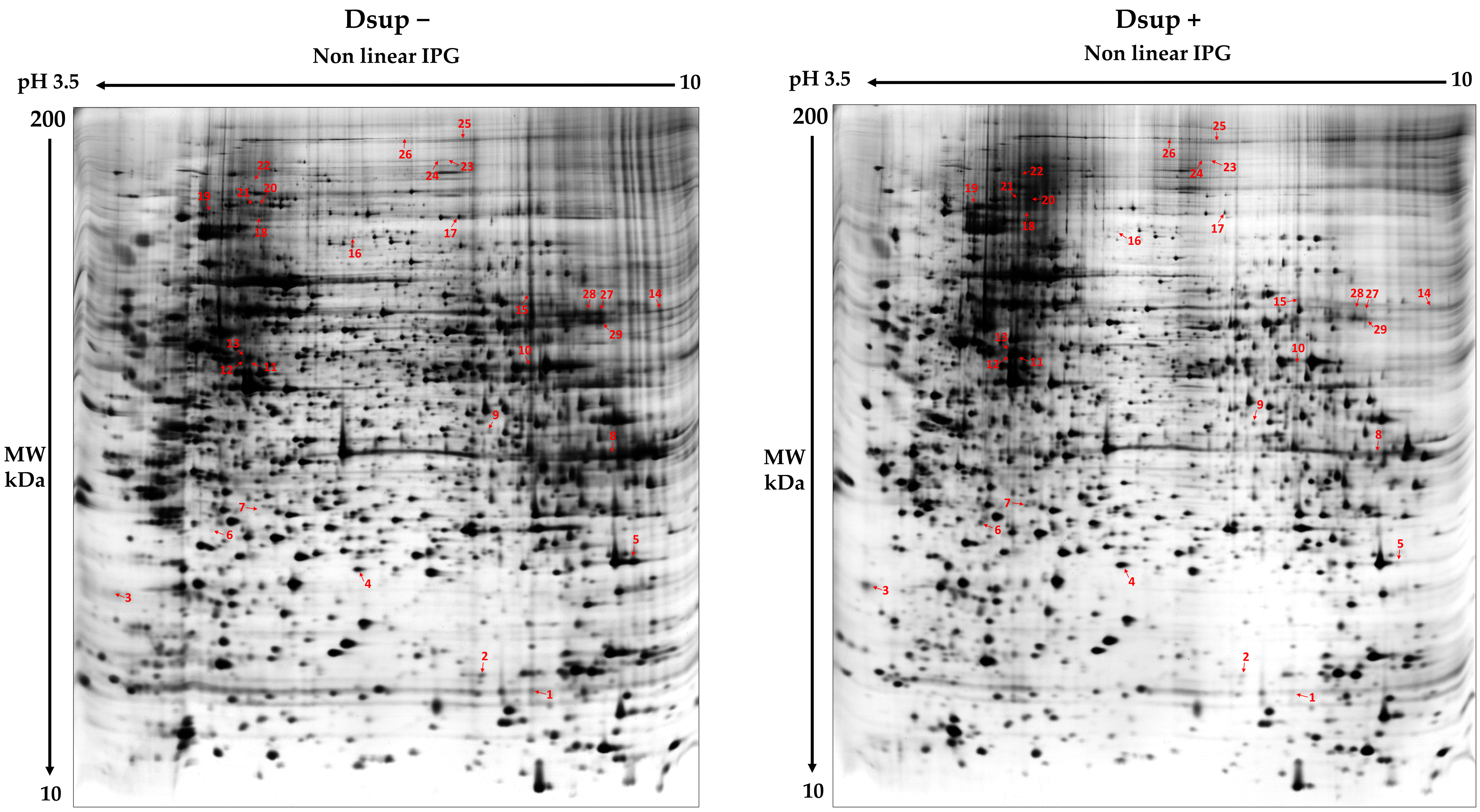


**Figure S1: Representative 2DE electropherograms of Dsup– vs Dsup+ HEK293T cells at basal condition.** The differentially abundant spots between the two considered conditions are reported in red. Spot numbers correspond to those showed in Table S1**.**

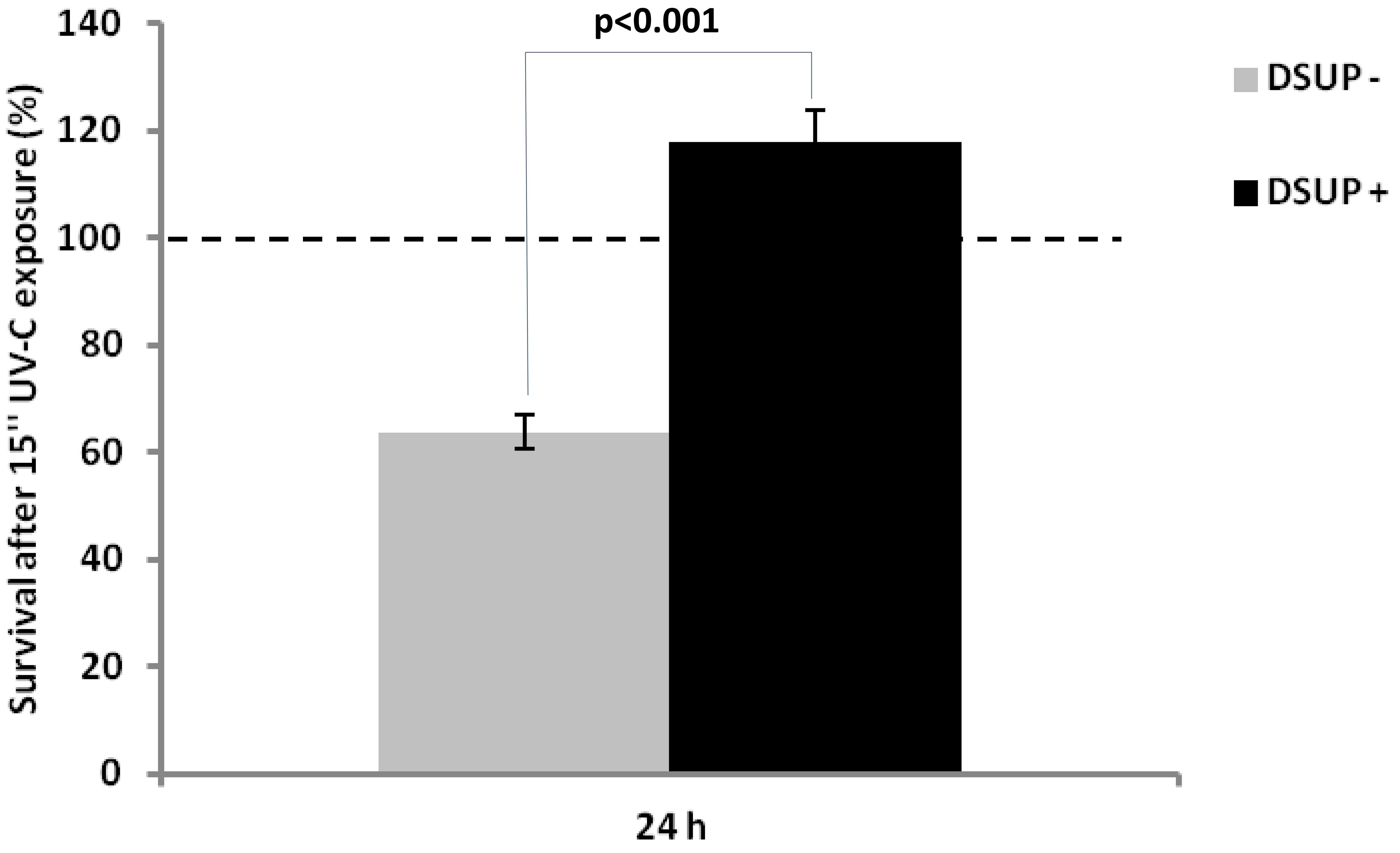


**Figure S2:** Survival assay of HEK293T cells control (Dsup-) and Dsup-transfected (Dsup+) after 15” of UV-C exposure recovered after 24h. The dotted line represents the basal condition (100%)

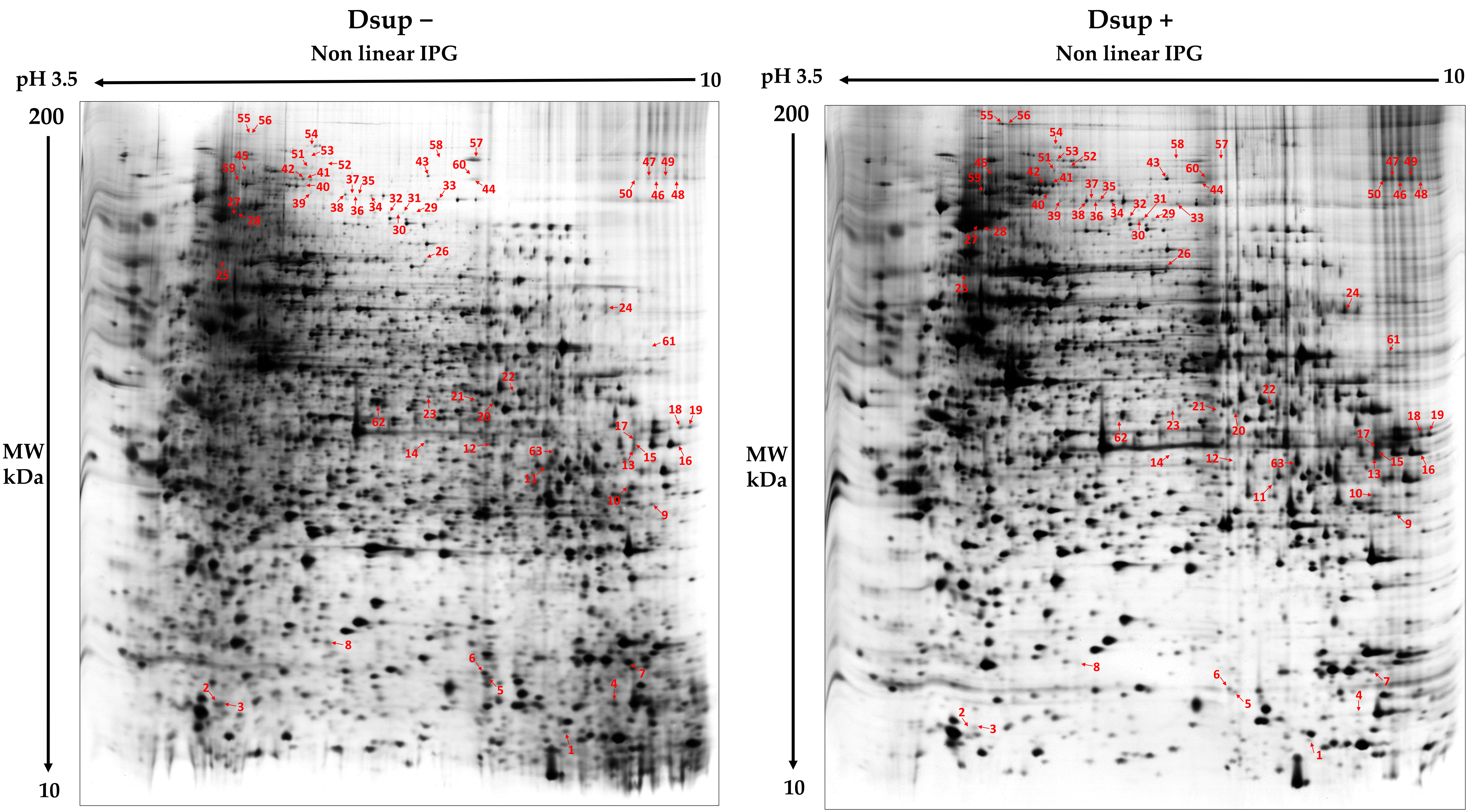


**Figure S3: Representative 2DE electropherograms of Dsup– vs Dsup+ HEK293T cells exposed to 15 seconds of UV-C radiation and recovered after 24h.** The differentially abundant spots between the two considered conditions are reported in red. Spot numbers correspond to that in Table S2.

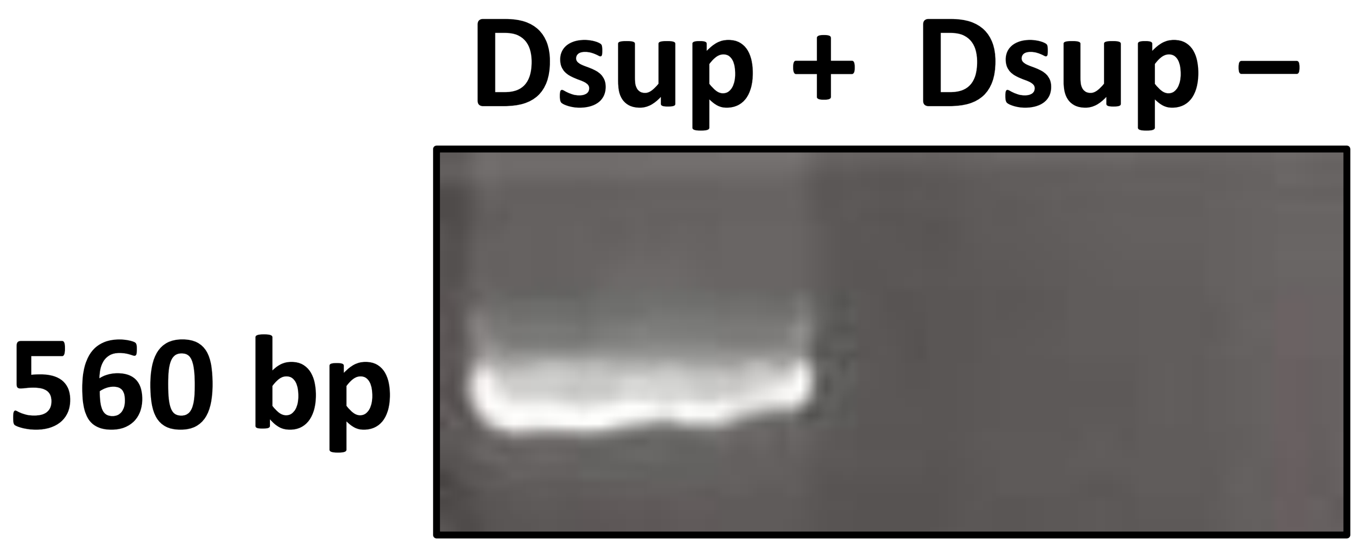


**Figure S4. Dsup expression assessment.** Dsup expression in control (Dsup−) and transfected (Dsup+) HEK293T cells was performed by endpoint PCR and visualized in 2% agarose gel by ethidium bromide.

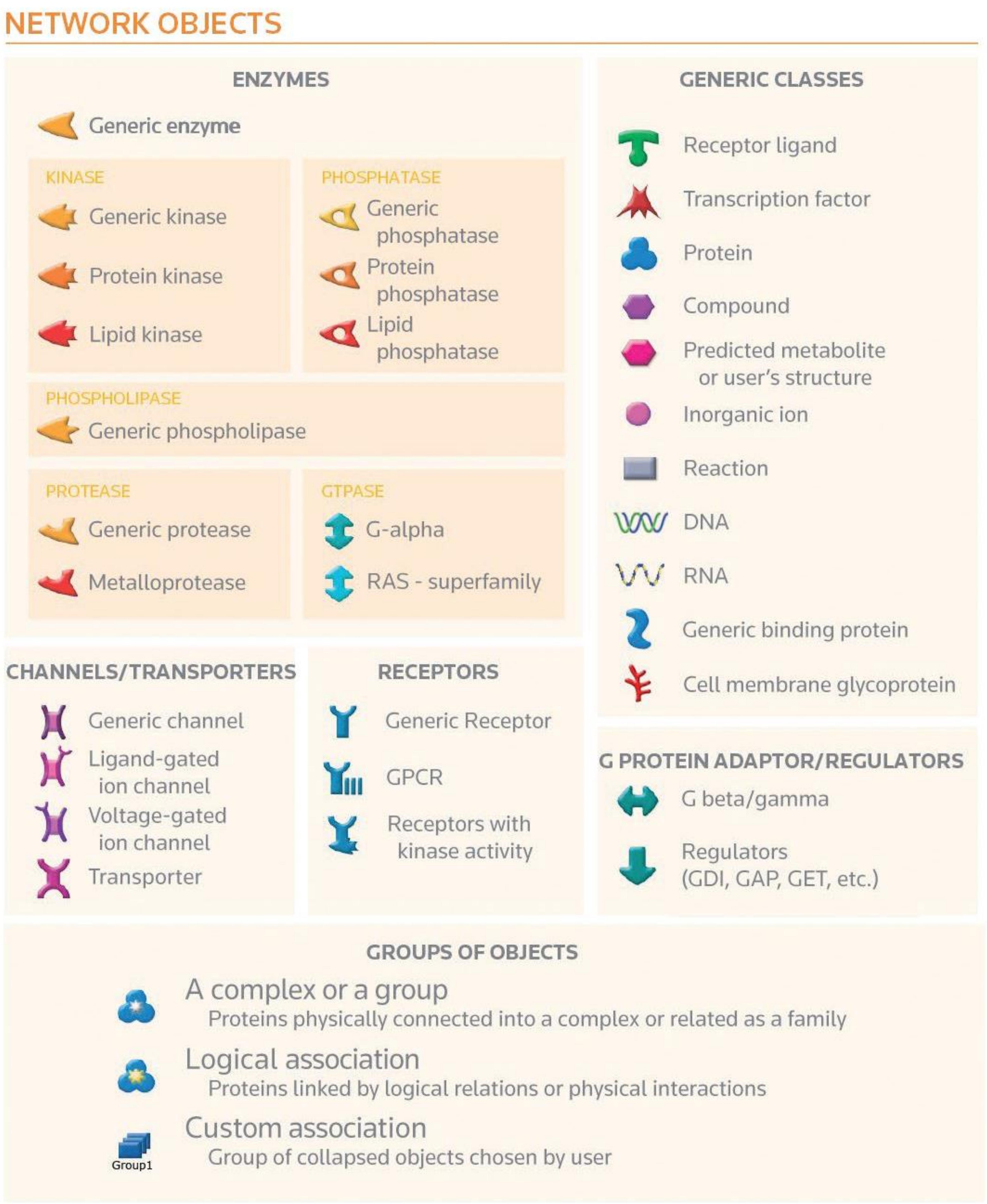


**Figure S5: Metacore legend**
