## supplementary tables for "Proteomics reveals how the tardigrade damage suppressor protein teaches transfected human cells to survive UV-C stress"

**Table S1. Proteomic data and identifications of differential spots in Dsup–/+ HEK293T cells at basal condition**

Table reports the spot number corresponding to that in Figure S3, protein name when the protein was identified by MALDI-ToF MS, UniProt Entry name and accession number (AC), the ANOVA Test and the mean of the %V of the specific spot in Dsup− and Dsup+ cells, the fold change of the %V means of Dsup− vs Dsup+ and viceversa, pI and MW. Last part of the table is dedicated to Mascot Search Results such as Score and Expect, Matched Peptides and sequence Coverage (%).

<table>
<tr><th rowspan="2">Spot n°</th><th rowspan="2">Protein name</th><th rowspan="2">Entry name</th><th rowspan="2">AC</th><th colspan="3">Anova test</th><th colspan="2">Ratio</th><th rowspan="2">pI - MW</th><th colspan="4">Mascot Search Results</th></tr>
<tr><th>Anova (p)</th><th>(Basal) Dsup−</th><th>(Basal) Dsup+</th><th>(Basal) Dsup−/+</th><th>(Basal) Dsup+/−</th><th colspan="2">Score<br>Expect</th><th>Matched peptides</th><th>Coverage (%)</th></tr>
<tr><td>1</td><td colspan="3"></td><td>0.003</td><td>0.043355</td><td>0.016045</td><td>2.702161872</td><td>0.370074054</td><td colspan="5"></td></tr>
<tr><td>2</td><td colspan="3"></td><td>0.011</td><td>0.019463</td><td>0.007949</td><td>2.448426838</td><td>0.408425518</td><td colspan="5"></td></tr>
<tr><td>3</td><td>Prostaglandin E synthase 3</td><td>TEBP_HUMAN</td><td>Q15185</td><td>0.040</td><td>0.025233</td><td>0.050755</td><td>0.497158563</td><td>2.011430708</td><td>4.35 - 18971</td><td colspan="2">128<br>3.2E$^{-09}$</td><td>8/12</td><td>45</td></tr>
<tr><td rowspan="2">4</td><td>Heat shock 70 kDa protein 1A N-term fragment</td><td>HS71A_HUMAN</td><td>P0DMV8</td><td rowspan="2">0.009</td><td rowspan="2">0.074455</td><td rowspan="2">0.165706</td><td rowspan="2">0.449321792</td><td rowspan="2">2.225576452</td><td>5.48 - 70294</td><td rowspan="2">224</td><td>169<br>2.6E$^{-13}$</td><td>17/30</td><td>26</td></tr>
<tr><td>Heat shock cognate 71 kDa protein N-term fragment</td><td>HSP7C_HUMAN</td><td>P11142</td><td>5.37 - 71082</td><td>89<br>2.3E$^{-05}$</td><td>12/30</td><td>21</td></tr>
<tr><td>5</td><td>Peroxiredoxin-1</td><td>PRDX1_HUMAN</td><td>Q06830</td><td>0.018</td><td>0.161855</td><td>0.040361</td><td>4.010171485</td><td>0.249365895</td><td>8.27 - 22324</td><td colspan="2">230<br>2E$^{-19}$</td><td>13/15</td><td>62</td></tr>
<tr><td>6</td><td>Heat shock 70 kDa protein 1A C-term fragment</td><td>HS71A_HUMAN</td><td>P0DMV8</td><td>0.007</td><td>0.006086</td><td>0.038048</td><td>0.159949258</td><td>6.251982753</td><td>5.48 - 70294</td><td colspan="2">111<br>1.6E$^{-07}$</td><td>10/14</td><td>16</td></tr>
<tr><td>7</td><td>Pyruvate kinase PKM N-term fragment</td><td>KPYM_HUMAN</td><td>P14618</td><td>0.010</td><td>0.002012</td><td>0.0199</td><td>0.101115909</td><td>9.889640595</td><td>7.96 - 58470</td><td colspan="2">152<br>1.3E$^{-11}$</td><td>12/20</td><td>26</td></tr>
<tr><td>8</td><td>Heterogeneous nuclear ribonucleoproteins A2/B1</td><td>ROA2_HUMAN</td><td>P22626</td><td>0.036</td><td>0.07028</td><td>0.030466</td><td>2.306830143</td><td>0.433495289</td><td>8-97 - 37464</td><td colspan="2">173<br>1E$^{-13}$</td><td>12/15</td><td>42</td></tr>
<tr><td>9</td><td>Pyruvate kinase PKM N-term fragment</td><td>KPYM_HUMAN</td><td>P14618</td><td>0.046</td><td>0.004739</td><td>0.030441</td><td>0.155674606</td><td>6.423655244</td><td>7.96 - 58470</td><td colspan="2">181<br>1.6E$^{-14}$</td><td>14/17</td><td>27</td></tr>
</table>

| | | | | | | | | | | | | |
|---|---|---|---|---|---|---|---|---|---|---|---|---|
| **10** | **Alpha-enolase** | **ENOA_HUMAN** | **P06733** | 0.037 | 0.018212 | 0.008153 | **2.23389549** | 0.447648516 | 7.01 - 47481 | **122** $1.3E^{-08}$ | 8/10 | 16 |
| **11** | **Actin. cytoplasmic 1 / Actin. cytoplasmic 2** | **ACTB_HUMAN / ACTG_HUMAN** | **P60709 / P63261** | 0.037 | 0.046185 | 0.10382 | 0.444861478 | **2.247890748** | 5.29 - 42052<br>5.31 - 42108 | **197** $4.1E^{-16}$ | 18/30 | 45 |
| **12** | **Eukaryotic initiation factor 4A-II** | **IF4A2_HUMAN** | **Q14240** | 0.012 | 0.043265 | 0.132976 | 0.325356519 | **3.073551449** | 5.33 - 46601 | **93** $9.3E^{-06}$ | 10/28 | 26 |
| **13** | **Eukaryotic initiation factor 4A-II** | **IF4A2_HUMAN** | **Q14240** | 0.005 | 0.020374 | 0.046276 | 0.440278618 | **2.27128904** | 5.33 - 46601 | **75** $6.4E^{-04}$ | 9/32 | 25 |
| **14** | | | | 0.004 | 0.051314 | 0.025473 | **2.014449267** | 0.496413594 | | | | |
| **15** | **Asparagine synthetase [glutamine-hydrolyzing]** | **ASNS_HUMAN** | **P08243** | 0.030 | 0.058045 | 0.027783 | **2.089232774** | 0.478644607 | 6.39 - 64899 | **139** $2.6E^{-10}$ | 9/10 | 14 |
| **16** | | | | 0.020 | 0.005422 | 0.002635 | **2.057749068** | 0.485967903 | | | | |
| **17** | | | | 0.049 | 0.055061 | 0.02651 | **2.076969696** | 0.481470674 | | | | |
| **18** | **Striatin-4** | **STRN4_HUMAN** | **Q9NRL3** | 0.047 | 0.008072 | 0.020211 | 0.399375914 | **2.503906631** | 5.21 - 81287 | **103** $1E^{-06}$ | 8/11 | 13 |
| **19** | **Heat shock protein HSP 90-beta** | **HS90B_HUMAN** | **P08238** | 0.020 | 0.013532 | 0.038488 | 0.351600345 | **2.844138278** | 4.97 - 83554 | **168**<br>**106** $5.1E^{-07}$ | 14/36 | 21 |
| | **Cleavage and polyadenylation specificity factor subunit 2** | **CPSF2_HUMAN** | **Q9P2I0** | | | | | | 4.98 - 89286 | **84** $7.7E^{-05}$ | 15/36 | 16 |
| **20** | **DNA replication licensing factor MCM6** | **MCM6_HUMAN** | **Q14566** | 0.049 | 0.014905 | 0.033412 | 0.446097602 | **2.241661905** | 5.29 - 93801 | **163** $1E^{-12}$ | 16/25 | 19 |
| **21** | **DNA replication licensing factor MCM6** | **MCM6_HUMAN** | **Q14566** | 0.043 | 0.020144 | 0.041181 | 0.489161616 | **2.044314124** | 5.29 - 93801 | **146** $5.1E^{-11}$ | 13/17 | 15 |
| **22** | | | | 0.027 | 0.006606 | 0.020521 | 0.321895365 | **3.106599559** | | | | |
| **23** | | | | 0.013 | 0.000986 | 0.002336 | 0.421981066 | **2.369774571** | | | | |
| **24** | | | | 0.012 | 0.001525 | 0.004027 | 0.378789183 | **2.639990909** | | | | |
| **25** | | | | 0.045 | 0.002205 | 0.004461 | 0.494212097 | **2.023422748** | | | | |

| | | | | | | | | | | | | |
|---|---|---|---|---|---|---|---|---|---|---|---|---|
| **26** | | | | 0.035 | 0.002161 | 0.00501 | 0.431435301 | **2.31784464** | | | | |
| **27** | **Tyrosine--tRNA ligase. cytoplasmic** | **SYYC_HUMAN** | **P54577** | 0.027 | 0.097784 | 0.036367 | **2.688779653** | 0.371915935 | 6.61 - 59448 | **118** $3.2E^{-08}$ | 9/13 | 25 |
| **28** | **Tyrosine--tRNA ligase. cytoplasmic** | **SYYC_HUMAN** | **P54577** | 0.027 | 0.183837 | 0.081087 | **2.267169064** | 0.441078707 | 6.61 - 59448 | **119** $2.6E^{-08}$ | 10/15 | 21 |
| **29** | **T-complex protein 1 subunit delta** | **TCPD_HUMAN** | **P50991** | 0.013 | 0.174077 | 0.066644 | **2.612048545** | 0.382841277 | 7.96 - 58401 | **161** $1.6E^{-12}$ | 13/16 | 27 |

**Table S2: Proteomic data and identifications of differential spots in Dsup–/+ HEK293T cells at 24h recovery after 15 seconds of UV-C radiation**

Table reports the spot number corresponding to that in Figure S4, protein name when the protein was identified by MALDI-ToF MS, UniProt Entry name and accession number (AC), the ANOVA Test and the mean of the %V of the specific spot in Dsup− and Dsup+ cells, the fold change of the %V means of Dsup− vs Dsup+ and viceversa, pI and MW. Last part of the table is dedicated to Mascot Search Results such as Score and Expect, Matched Peptides and sequence Coverage (%).

| Spot n° | Protein name | Entry name | AC | Anova test | | | Ratio | | pI - MW | Mascot Search Results | | | |
|---|---|---|---|---|---|---|---|---|---|---|---|---|---|
| | | | | Anova (p) | (UV) Dsup− | (UV) Dsup+ | (UV) Dsup −/+ | (UV) Dsup +/− | | Score Expect | | Matched peptides | Coverage (%) |
| **1** | | | | 0.030 | 0.021045 | 0.0093442 | **2.252196461** | 0.444010999 | | | | | |
| **2** | **Regulator complex protein LAMTOR2** | **LTOR2_HUMAN** | **Q9Y2Q5** | 0.026 | 0.0190797 | 0.0040741 | **4.683227283** | 0.213527967 | 5.30 - 13613 | **103** | **78** 2.9E$^{-04}$ | 5/14 | 38 |
| | **Replication protein A 14 kDa subunit** | **RFA3_HUMAN** | **P35244** | | | | | | 4.96 - 13674 | | **63** 9.5E$^{-03}$ | 4/14 | 54 |
| **3** | **Thioredoxin** | **THIO_HUMAN** | **P10599** | 0.013 | 0.0270251 | 0.0090355 | **2.99097785** | 0.334338818 | 4.82 - 12015 | **69** 2.8E$^{-03}$ | | 5/7 | 32 |
| **4** | | | | 0.050 | 0.0481777 | 0.011223 | **4.292765278** | 0.232950077 | | | | | |
| **5** | | | | 0.005 | 0.0467749 | 0.0196331 | **2.382453412** | 0.419735385 | | | | | |
| **6** | | | | 0.028 | 0.0379288 | 0.0156038 | **2.430736037** | 0.411398023 | | | | | |
| **7** | **Peptidyl-prolyl cis-trans isomerase A** | **PPIA_HUMAN** | **P62937** | 0.015 | 0.0833672 | 0.0167316 | **4.982629633** | 0.200697237 | 7.68-18229 | **182** 1.3E$^{-14}$ | | 13/26 | 58 |
| **8** | **Actin-related protein 2/3 complex subunit 5** | **ARPC5_HUMAN** | **O15511** | 0.030 | 0.0273185 | 0.0094444 | **2.892558075** | 0.345714753 | 5.47-16367 | **146** 5.1E$^{-11}$ | | 11/17 | 52 |
| **9** | **Peroxiredoxin-1** | **PRDX1_HUMAN** | **Q06830** | 0.022 | 0.0184992 | 0.0615035 | 0.300783424 | **3.324651292** | 8.27- 22324 | **100** 2E$^{-06}$ | | 7/16 | 35 |
| **10** | | | | 0.042 | 0.0711136 | 0.0263909 | **2.694627939** | 0.371108748 | | | | | |
| **11** | **S-** | **ESTD_HUMAN** | **P10768** | 0.037 | 0.0474349 | 0.0213141 | **2.225519297** | 0.449333331 | 6.54-31956 | **158** | | 11/17 | 50 |

| | | | | | | | | | | | | | |
|---|---|---|---|---|---|---|---|---|---|---|---|---|---|
| | **formylglutathione hydrolase** | | | | | | | | | | $3.2E^{-12}$ | | |
| **12** | **Phosphatidylinositol transfer protein beta isoform** | **PIPNB_HUMAN** | **P48739** | 0.010 | 0.0342307 | 0.0064848 | **5.278594214** | 0.189444378 | 6.41-31805 | | **356** $5.1E^{-32}$ | 23/26 | 67 |
| **13** | **Heterogeneous nuclear ribonucleoproteins A2/B1** | **ROA2_HUMAN** | **P22626** | 0.010 | 0.0270482 | 0.0973908 | 0.277728245 | **3.600642064** | 8.97-37464 | | **143** $1E^{-10}$ | 10/15 | 38 |
| **14** | | | | 0.049 | 0.0118442 | 0.0021141 | **5.602421492** | 0.178494246 | | | | | |
| **15** | **Heterogeneous nuclear ribonucleoproteins A2/B1** | **ROA2_HUMAN** | **P22626** | 0.001 | 0.0135462 | 0.0539949 | 0.250879095 | **3.985983769** | 8.97-37464 | | **213** $1E^{-17}$ | 14/17 | 42 |
| **16** | **Heterogeneous nuclear ribonucleoproteins A2/B1** | **ROA2_HUMAN** | **P22626** | 0.008 | 0.0281454 | 0.0620498 | 0.453592945 | **2.204619825** | 8.97-37464 | | **114** $8.1E^{-08}$ | 8/13 | 35 |
| **17** | **Heterogeneous nuclear ribonucleoproteins A2/B1** | **ROA2_HUMAN** | **P22626** | 0.042 | 0.0395889 | 0.1261191 | 0.313901255 | **3.1857152** | 8.97-37464 | | **186** $5.1E^{-15}$ | 13/18 | 42 |
| **18** | **Heterogeneous nuclear ribonucleoproteins A2/B1** | **ROA2_HUMAN** | **P22626** | <0.0001 | 0.0243909 | 0.064495 | 0.378182093 | **2.644228851** | 8.97-37464 | | **344** $8.1E^{-31}$ | 23/25 | 53 |
| **19** | **Heterogeneous nuclear ribonucleoproteins A2/B1** | **ROA2_HUMAN** | **P22626** | <0.0001 | 0.0170888 | 0.0497053 | 0.343801701 | **2.908653443** | 8.97-37464 | | **167** $4.1E^{-13}$ | 11/14 | 42 |
| **20** | **Poly(rC)-binding protein 1** | **PCBP1_HUMAN** | **Q15365** | 0.035 | 0.0688577 | 0.0337199 | **2.042048324** | 0.489704376 | 6.66-37987 | | **137** $4.1E^{-10}$ | 10/15 | 41 |
| **21** | | | | 0.016 | 0.0061338 | 0.0013465 | **4.555468387** | 0.219516395 | | | | | |
| **22** | **Mitogen-activated protein kinase** | **MK01_HUMAN** | **P28482** | 0.019 | 0.0164557 | 0.0342318 | 0.480713334 | **2.080241862** | 6.50 - 41762 | **175** | **126** $5.1e^{-09}$ | 10/20 | 24 |

| | | | | | | | | | | | | |
|---|---|---|---|---|---|---|---|---|---|---|---|---|
| | **1** | | | | | | | | | | | |
| | **Sialic acid synthase** | **SIAS_HUMAN** | **Q9NR45** | | | | | | 6.29 - 40738 | **90**<br>2.2E$^{-05}$ | 8/20 | 23 |
| **23** | **Translation initiation factor eIF-2B subunit beta** | **EI2BB_HUMAN** | **P49770** | 0.047 | 0.0113539 | 0.0031515 | **3.602685479** | 0.277570719 | 5.77-39193 | **216**<br>5.1E$^{-18}$ | 17/24 | 45 |
| **24** | **D-3-phosphoglycerate dehydrogenase** | **SERA_HUMAN** | **O43175** | 0.003 | 0.022067 | 0.07731 | 0.285434953 | **3.50342518** | 6.29-57356 | **160**<br>2E$^{-12}$ | 14/24 | 31 |
| **25** | **Heat shock 70 kDa protein 1A** | **HS71A_HUMAN** | **P0DMV8** | 0.037 | 0.03986 | 0.0879722 | 0.453097683 | **2.207029604** | 5.48- 70294 | **275**<br>6.4E$^{-24}$ | 21/29 | 43 |
| | **Heat shock 70 kDa protein 1B** | **HS71B_HUMAN** | **P0DMV9** | | | | | | 5.48- 70294 | **275**<br>6.4E$^{-24}$ | 21/29 | 43 |
| **26** | **Heat shock 70 kDa protein 1A** | **HS71A_HUMAN** | **P0DMV8** | 0.047 | 0.0059819 | 0.0130175 | 0.459529213 | **2.176140214** | 5.48- 70294 | **161**<br>1.6E$^{-12}$ | 12/15 | 23 |
| | **Heat shock 70 kDa protein 1B** | **HS71B_HUMAN** | **P0DMV9** | | | | | | 5.48- 70294 | **161**<br>1.6E$^{-12}$ | 12/15 | 23 |
| **27** | **Heat shock protein HSP 90-beta** | **HS90B_HUMAN** | **P08238** | 0.017 | 0.0833278 | 0.1669537 | 0.499107138 | **2.003577839** | 4.97-83554 | **169**<br>2.6E$^{-13}$ | 20/35 | 29 |
| **28** | **Heat shock protein HSP 90-beta** | **HS90B_HUMAN** | **P08238** | 0.032 | 0.040419 | 0.0918989 | 0.439820484 | **2.273654904** | 4.97-83554 | **169**<br>2.6E$^{-13}$ | 20/35 | 29 |
| **29** | **Cold shock domain-containing protein E1** | **CSDE1_HUMAN** | **O75534** | 0.048 | 0.003588 | 0.0085877 | 0.417811 | **2.393426693** | 5.88-89684 | **117**<br>4.1E$^{-08}$ | 11/17 | 17 |
| **30** | **Cold shock domain-containing protein E1** | **CSDE1_HUMAN** | **O75534** | 0.003 | 0.0022853 | 0.0064755 | 0.352919859 | **2.833504472** | 5.88-89684 | **236**<br>5.1E$^{-20}$ | 21/28 | 28 |
| **31** | **Cold shock domain-** | **CSDE1_HUMAN** | **O75534** | 0.047 | 0.0035469 | 0.0075604 | 0.469145453 | **2.131535099** | 5.88-89684 | **101**<br>1.6E$^{-06}$ | 9/13 | 13 |

| | | | | | | | | | | | | |
|---|---|---|---|---|---|---|---|---|---|---|---|---|
| | **containing protein E1** | | | | | | | | | | | |
| **32** | **Cold shock domain-containing protein E1** | **CSDE1_HUMAN** | **O75534** | 0.048 | 0.0046699 | 0.0097517 | 0.478884216 | **2.088187429** | 5.88-89684 | **74**<br>$7.9E^{-04}$ | 8/18 | 14 |
| **33** | **Elongation factor 2** | **EF2_HUMAN** | **P13639** | 0.032 | 0.0081677 | 0.0196089 | 0.416530617 | **2.400783901** | 6.41-96246 | **178**<br>$3.2E^{-14}$ | 16/20 | 21 |
| **34** | **Neutral alpha-glucosidase AB** | **GANAB_HUMAN** | **Q14697** | 0.028 | 0.0068976 | 0.0216881 | 0.318036493 | **3.144293254** | 5.74-107263 | **316**<br>$5.1E^{-28}$ | 29/34 | 32 |
| **35** | **Heat shock 70 kDa protein 4L** | **HS74L_HUMAN** | **O95757** | 0.016 | 0.0040901 | 0.0144419 | 0.283208476 | **3.530967761** | 5.63 - 95479 | **61**<br>$1.5E^{-02}$ | 5/6 | 8 |
| **36** | **Heat shock 70 kDa protein 4L** | **HS74L_HUMAN** | **O95757** | 0.028 | 0.0016363 | 0.0072614 | 0.225338917 | **4.437759864** | 5.63-95479 | **124**<br>$8.1E^{-09}$ | 13/20 | 16 |
| **37** | **Heat shock 70 kDa protein 4L** | **HS74L_HUMAN** | **O95757** | 0.021 | 0.0012532 | 0.0049527 | 0.253033774 | **3.952041592** | 5.63-95479 | **124**<br>$8.1E^{-09}$ | 13/20 | 16 |
| **38** | **Neutral alpha-glucosidase AB** | **GANAB_HUMAN** | **Q14697** | 0.021 | 0.0079115 | 0.0167872 | 0.471280358 | **2.121879224** | 5.74-107263 | **97**<br>$2.9E^{-06}$ | 10/17 | 12 |
| **39** | **Ubiquitin-like modifier-activating enzyme 1** | **UBA1_HUMAN** | **P22314** | 0.015 | 0.0010479 | 0.0048447 | 0.216293867 | **4.623339591** | 5.49 - 118858 | **57**<br>$3.9E^{-02}$ | 4/4 | 6 |
| **40** | **Ubiquitin-like modifier-activating enzyme 1** | **UBA1_HUMAN** | **P22314** | 0.019 | 0.0099294 | 0.0352576 | 0.281624783 | **3.550823865** | 5.49- 118858 | **307**<br>$4.1E^{-27}$ | 25/30 | 35 |
| **41** | **Ubiquitin-like modifier-activating enzyme 1** | **UBA1_HUMAN** | **P22314** | 0.008 | 0.0075397 | 0.033307 | 0.226370933 | **4.417528287** | 5.49- 118858 | **165**<br>$6.4E^{-13}$ | 17/29 | 24 |
| **42** | **Ubiquitin-like modifier-activating enzyme 1** | **UBA1_HUMAN** | **P22314** | 0.001 | 0.0120293 | 0.0314904 | 0.381999189 | **2.617806604** | 5.49- 118858 | **303**<br>$1E^{-26}$ | 25/31 | 36 |
| **43** | **Transcription intermediary factor 1-beta** | **TIF1B_HUMAN** | **Q13263** | 0.018 | 0.0103278 | 0.0218028 | 0.473691565 | **2.111078334** | 5.52-90261 | **77**<br>$3.7E^{-04}$ | 7/11 | 10 |
| **44** | | | | 0.010 | 0.0121098 | 0.0732296 | 0.165366965 | **6.047157017** | | | | |

| | | | | | | | | | | | | |
|---|---|---|---|---|---|---|---|---|---|---|---|---|
| **45** | **Transcription intermediary factor 1-beta** | **TIF1B_HUMAN** | **Q13263** | 0.005 | 0.0151401 | 0.0435923 | 0.347311072 | **2.879263237** | 5.52-90261 | **109** 2.6E$^{-07}$ | 11/22 | 16 |
| **46** | | | | 0.002 | 0.0122145 | 0.0421808 | 0.289574878 | **3.453338246** | | | | |
| **47** | | | | 0.027 | 0.0190551 | 0.0476007 | 0.40031134 | **2.498055639** | | | | |
| **48** | **Heterogeneous nuclear ribonucleoprotein U** | **HNRPU_HUMAN** | **Q00839** | 0.004 | 0.0217492 | 0.0640288 | 0.339679092 | **2.943955113** | 5.76-91269 | **60** 8.5E$^{-04}$ | 7/14 | 10 |
| **49** | **Heterogeneous nuclear ribonucleoprotein U** | **HNRPU_HUMAN** | **Q00839** | 0.000 | 0.0198534 | 0.0624223 | 0.318049435 | **3.144165305** | 5.76-91269 | **110** 2E$^{-07}$ | 8/9 | 13 |
| **50** | **Heterogeneous nuclear ribonucleoprotein U** | **HNRPU_HUMAN** | **Q00839** | 0.027 | 0.0166104 | 0.043295 | 0.383655541 | **2.606504773** | 5.76-91269 | **70** 1.9E$^{-03}$ | 5/5 | 9 |
| **51** | **DNA replication licensing factor MCM2** | **MCM2_HUMAN** | **P49736** | 0.029 | 0.0012638 | 0.0041363 | 0.305540499 | **3.272888545** | 5.34-102516 | **165** 6.4E$^{-13}$ | 17/27 | 25 |
| **52** | | | | 0.039 | 0.0012606 | 0.0041951 | 0.300503048 | **3.327753268** | | | | |
| **53** | **Hypoxia up-regulated protein 1** | **HYOU1_HUMAN** | **Q9Y4L1** | 0.029 | 0.0021725 | 0.0067587 | 0.321432887 | **3.111069342** | 5.16-111494 | **101** 1.6E$^{-06}$ | 8/8 | 10 |
| **54** | | | | 0.039 | 0.0014627 | 0.0058263 | 0.25104775 | **3.983305971** | | | | |
| **55** | | | | 0.015 | 0.0065333 | 0.0216408 | 0.301897653 | **3.312380835** | | | | |
| **56** | | | | 0.009 | 0.0043338 | 0.0135865 | 0.318976634 | **3.135025874** | | | | |
| **57** | | | | 0.001 | 0.0459592 | 0.0153109 | **3.001739505** | 0.333140167 | | | | |
| **58** | | | | 0.048 | 0.001202 | 0.0048902 | 0.245797664 | **4.068386919** | | | | |
| **59** | **Eukaryotic translation initiation factor 3 subunit B** | **EIF3B_HUMAN** | **P55884** | 0.026 | 0.0161619 | 0.0323538 | 0.499537406 | **2.001852089** | 4.89-92823 | **153** 1E$^{-11}$ | 14/18 | 20 |
| **60** | | | | 0.040 | 0.0165049 | 0.0848576 | 0.194500986 | **5.141362117** | | | | |
| **61** | **Alpha-enolase** | **ENOA_HUMAN** | **P06733** | 0.007 | 0.0073665 | 0.0307339 | 0.23968621 | **4.172121541** | 7.01 - 47481 | **197** | 15/20 | 32 |

| | | | | | | | | | | | | |
|---|---|---|---|---|---|---|---|---|---|---|---|---|
| | | | | | | | | | | 4.1E-16 | | |
| **62** | **Isocitrate dehydrogenase [NAD] subunit alpha. mitochondrial** | **IDH3A_HUMAN** | **P50213** | 0.004 | 0.0408057 | 0.0139044 | **2.934736121** | 0.340746138 | 6.47 - 40022 | **93** 9.8E-06 | 7/12 | 19 |
| **63** | **Ribose-phosphate pyrophosphokinase 1** | **PRPS1_HUMAN** | **P60891** | 0.001 | 0.0350293 | 0.0925712 | 0.378404227 | **2.642676614** | 6.51-35325 | **152** 1.3E-11 | 10/14 | 35 |
